# Combinatorial Control of Corticospinal Axon Growth by Retinoic Acid Receptors

**DOI:** 10.64898/2026.07.24.740479

**Authors:** Yogesh Sahu, Athul PS Narayan, Manojkumar Kumaran, Shringika Soni, Prakash Chermakani, Dhruvakumar Kesireddy, Meghana Konda, Anisha S Menon, Ishwariya Venkatesh

## Abstract

Regeneration of central nervous system (CNS) axons depends on Transcription Factors (TFs) that reactivate developmental growth programs, yet most such factors remain unknown. By intersecting developmental chromatin binding with pro-growth gene networks, we identified two retinoic acid receptor transcription factors, RARA and RARG, whose occupancy at growth-associated genes is progressively lost as neurons mature. Restoring both factors together increased neurite outgrowth beyond either alone in two independent systems, the Neuro-2a cell line and primary cortical neurons. *In vivo*, the same combination drove cross-midline sprouting after pyramidotomy and long-tract regeneration after thoracic spinal cord crush, with concordant recovery of hindlimb gait and grip strength. Interestingly, neither receptor alone was sufficient, hinting at combinatorial regulation. Single-nucleus transcriptomics delineated that only the combination reactivated relevant cytoskeletal and gene-expression programs, while genome-wide binding maps showed that RARA and RARG partition the regulatory landscape, with RARG dominating promoters and RARA occupying distal enhancers, so that neither receptor reconstitutes the developmental growth state alone. Intriguingly, this cooperative requirement was specific to the CNS: in peripheral sensory neurons, RARG alone was sufficient and RARA was dispensable. These data identify RARA and RARG as novel cooperative regulators of regenerative axon growth in mammalian CNS and PNS neurons and potential targets for therapeutic intervention.

## Introduction

After central nervous system (CNS) injury, severed axons fail to regrow, and this failure is driven in large part by a decline in the intrinsic growth capacity of neurons as they mature^1,2^. One strategy to overcome this is to reintroduce the transcription factors that specify axon growth during development, reactivating a growth program that adult neurons have switched off. Several such factors have been identified, including KLF family members, SOX11, and others, and forced expression of individual factors can promote neurite outgrowth and, in some cases, axon regeneration after injury^3–5^. Yet the set of transcription factors capable of driving CNS regeneration is far from complete, and single factors typically produce partial effects, pointing to regulators and combinations that have not yet been tested.

Retinoic acid signaling is one candidate axis. Retinoic acid receptors are ligand-activated nuclear transcription factors, and among them RARβ has been the main focus of regeneration studies, where RARβ2 expression promotes neurite outgrowth and axon regrowth after spinal cord injury^6–9^. The other two isoforms, RARα (RARA) and RARγ (RARG), are also expressed in neurons and regulate developmental gene programs^10,11^, but their contribution to adult CNS axon growth is largely untested, and whether they act together has not been examined at all. Because nuclear receptors frequently function as dimers and can regulate distinct target sets, the effect of expressing RARA and RARG in combination cannot be predicted from either factor alone, leaving a specific and testable gap.

Here we show that RARA and RARG cooperate to drive axon growth in the CNS. Their binding to a pro-growth gene network is highest in the embryonic brain and declines with maturation, mirroring the loss of intrinsic growth capacity. Neither receptor is sufficient on its own: combined expression is required to increase neurite outgrowth in cultured neurons, to drive corticospinal sprouting after pyramidotomy, and to support axon regeneration and functional recovery after thoracic spinal cord injury. Single-nucleus RNA sequencing and CUT&RUN show why co-delivery is needed, as the two receptors occupy largely separate genomic compartments and, together, regulate a combination-specific set of target genes that neither factor engages alone. This cooperative requirement is specific to the CNS: in peripheral sensory neurons, RARG alone is sufficient and RARA is dispensable. RARA and RARG therefore act as a combinatorial switch for corticospinal growth, and the requirement for both is a central, rather than universal, feature of the RAR growth program.

## Results

### In silico analysis reveals dynamic binding of retinoic acid receptors to a pro-growth gene network across development

To probe how RARA and RARG might act in CNS development and axon regeneration, we combined age-matched ATAC-seq datasets from ENCODE with Gene Ontology (GO) analysis across three mouse developmental stages: embryonic, postnatal, and adult.

To curate a growth-associated gene network, literature text mining for neural repair and development terms identified 2,172 candidate genes. Subsequent Gene Ontology and canonical pathway filtering for axonal dynamics and neural repair yielded a standardized network of 628 unique genes (Supplementary table S1). GO enrichment of this gene set confirmed its functional coherence, with terms clustering around axon guidance, neurogenesis, and developmental growth (Fig. 1a) (Supplementary table S1). To test whether preferential binding to the pro-growth network is specific to the retinoic acid receptors, we included one positive and two negative control transcription factors in the same analysis (Fig. 1b) (Supplementary table S1): KLF6, an established pro-growth factor, and TBR1 and BCL11B, two cortical factors, but not expected to target pro-growth genes. KLF6 bound the largest fraction of pro-growth genes (∼69%), while the negative controls TBR1 (∼18%) and BCL11B (∼27%) bound far fewer. RARA and RARG bound well above both negative controls and approached the KLF6 benchmark, placing their occupancy within the range expected of genuine pro-growth regulators rather than factors unrelated to axon growth.

**Figure 1:**
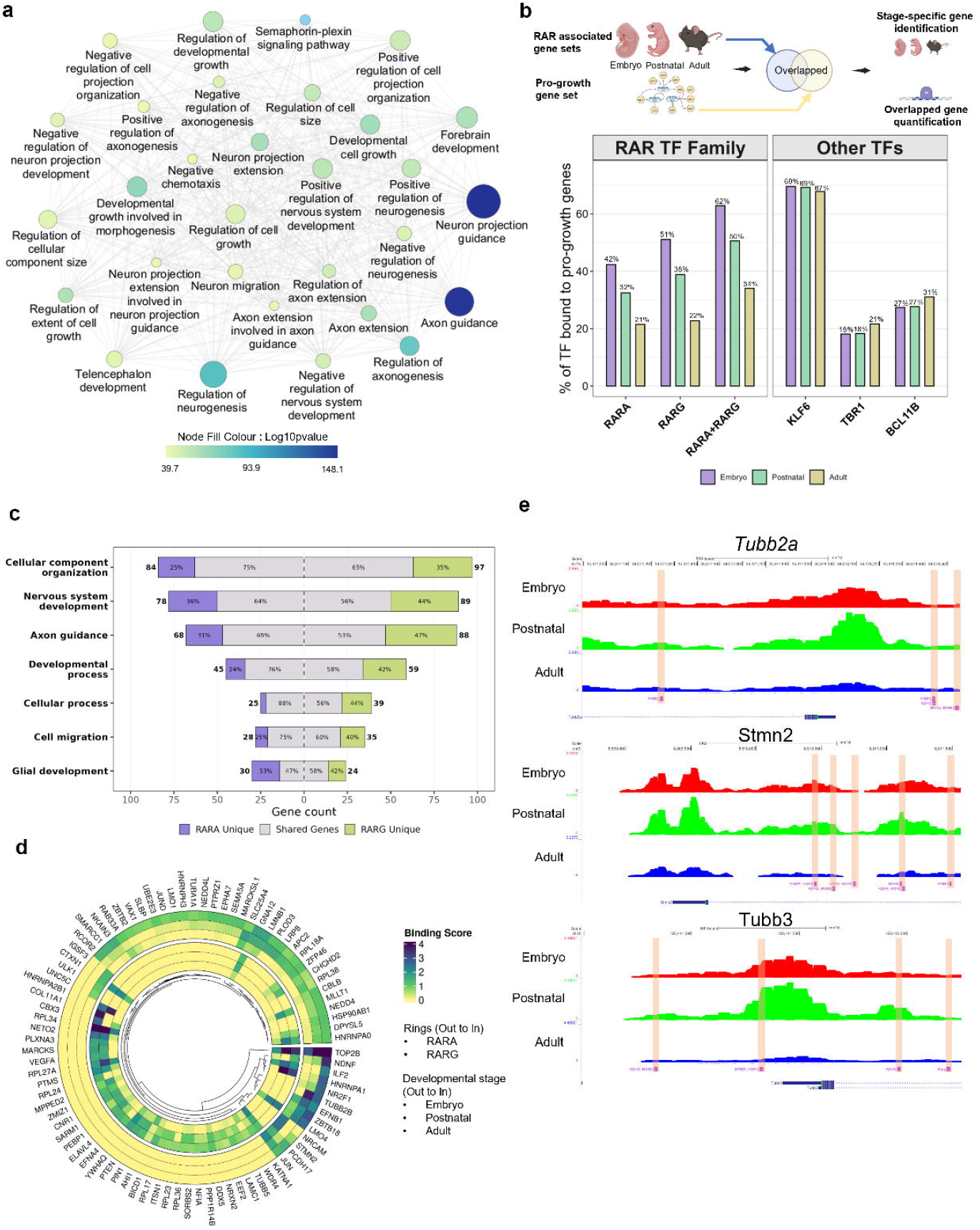
Developmental binding dynamics and functional enrichment of Retinoic Acid Receptors (RARA and RARG) within pro-growth gene networks. **a.** Gene Ontology (GO) network of pro-growth genes. Cytoscape network visualization depicting the enriched biological processes and developmental pathways associated with a curated list of 628 unique pro-growth genes. Nodes represent individual GO terms, with edges indicating interconnectivity between related developmental, morphological, and neurogenic pathways. **b.** Transcription factor binding prevalence across developmental stages. *Top:* Schematic illustrating the computational workflow used to identify the intersection between RAR-associated gene sets (across Embryo, Postnatal, and Adult stages) and the 628 pro-growth gene list. *Bottom:* Bar plot quantifying the percentage of pro-growth genes bound by RARA, RARG, and their combination (RARA+RARG), alongside control transcription factors (KLF6, TBR1, and BCL11B). Data is grouped by developmental stage: Embryo (purple), Postnatal (green), and Adult (yellow). **c.** Differential GO enrichment by RAR isoforms. Diverging tornado plot highlighting key biological processes enriched by RARA and RARG binding. The plot displays the total gene count per GO term, segmented by the proportion (%) of genes uniquely bound by RARA (purple), uniquely bound by RARG (green), or shared between both isoforms (grey). **d.** Binding affinity of RARs to highly variant genes. Circular heatmap representing the binding scores of RARA and RARG to the top 50 highly variant genes across developmental epochs. Concentric rings (from outer to inner) represent the binding profiles for RARA followed by RARG. Within each transcription factor block, the constituent rings (from outer to inner) denote the Embryo, Postnatal, and Adult stages, respectively. Colour intensity (yellow to dark blue) corresponds to the relative binding score. **e.** Chromatin accessibility at key regeneration-associated loci. Genome browser snapshots displaying representative binding sites and chromatin accessibility profiles at the *Tubb2a*, *Stmn2*, and *Tubb3* loci. Tracks visualize ENCODE ATAC-seq data from Embryo (red), Postnatal (green), and Adult (blue) stages, highlighting dynamic developmental shifts in accessibility at these critical pro-growth genes. Vertical shaded bands indicate specific regions of differential binding interest.

To separate the shared and distinct roles of the two receptors, we mapped RARA-and RARG-bound genes onto the top-enriched GO terms (Fig. 1c) (Supplementary table S1). Both targeted gene networks tied to neural repair, with the largest gene counts in cellular component organization (RARA-84, RARG-97), nervous system development (RARA-78, RARG-89), and axon guidance (RARA-68, RARG-88). Most terms showed heavy overlap: in cellular component organization and developmental process, 75% and 76% of RARA-bound genes, respectively, were also bound by RARG. Beyond this shared core, RARG regulated both more genes and a larger fraction of unique targets than RARA in most categories, most clearly in axon guidance (47% of RARG targets unique vs. 31% for RARA) and nervous system development (44% vs. 36%). Glial development ran the other way: RARA bound more genes (30 vs. 24) and held a higher unique fraction (53%), pointing to a distinct RARA-driven program in glial pathways. We then visualized RARA and RARG binding scores across the top 50 most variable genes as a circular heatmap spanning the three stages (Fig. 1d) (Supplementary table S1). For both receptors, binding was strongest in the embryo and weakened through the postnatal and adult stages, so the genome-wide decline also held at the level of individual high-variance targets.

Genome browser tracks at three hallmark growth genes such as *tubb2a*, *stmn2*, and *tubb3* showed the same binding pattern locus by locus (Fig. 1e). Both receptors gave clear binding peaks in the embryonic and postnatal cortex, and signal fell sharply in the adult tracks. RARA and RARG occupancy at pro-growth loci is therefore an early-developmental feature that the mature CNS largely loses.

### RARA and RARG synergistically enhance neurite outgrowth in cultured neurons

Building on the *in silico* prioritization of RARA and RARG as candidate regulators of neurite growth, we asked whether overexpressing them could drive the phenotype directly. We used AAV vectors encoding each receptor to transduce Neuro-2a (N2A) cells as an initial screening platform (Fig. 2a–c). Cells received AAV-tdTOMATO (control), AAV-RARG, AAV-RARA, or both receptors together, then were switched to differentiation medium, fixed, and immunostained for βIII-tubulin for neurite tracing and quantification. Mean neurite length rose across groups, from 101.3 µm in AAV-tdTOMATO controls to 145.8 µm with AAV-RARG and 163.8 µm with AAV-RARA, and reached 203.3 µm with combined AAV-RARA and AAV-RARG. One-way ANOVA confirmed a significant effect of treatment (F(3,1369) = 109.7, p < 0.0001). By Tukey’s post hoc test, every treatment group differed from control (p < 0.0001 in each case), and the combination outperformed either single factor (p < 0.0001 vs. AAV-RARA; p < 0.0001 vs. AAV-RARG). The two single-factor groups differed only marginally (AAV-RARA vs. AAV-RARG, p = 0.0498) and behaved as comparable individual drivers in this assay (Supplementary table S2). To test whether this activity extends beyond an immortalized line, we repeated the experiment in primary cortical neurons isolated from post-natal day 0 mouse brain (Fig. 2d–f). Neurons were transduced with AAV-GFP (control) or with combined AAV-RARA and AAV-RARG, kept for two days *in vitro*, and stained for βIII-tubulin prior to quantification. Combined receptor expression again increased neurite length relative to control (283.97 ± 122.0 µm, n = 65 neurites, versus 86.99 ± 39.9 µm, n = 55 neurites for AAV-GFP), a 3.3-fold increase. The difference was significant by Mann-Whitney U test (p < 0.0001, exact) (Supplementary table S3). All the virus capsid and integrity were checked using SDS-PAGE (Supplementary figure 1). Overexpression of RARA and RARG viral constructs were confirmed using western blot (Supplementary figure 2).

**Figure 2:**
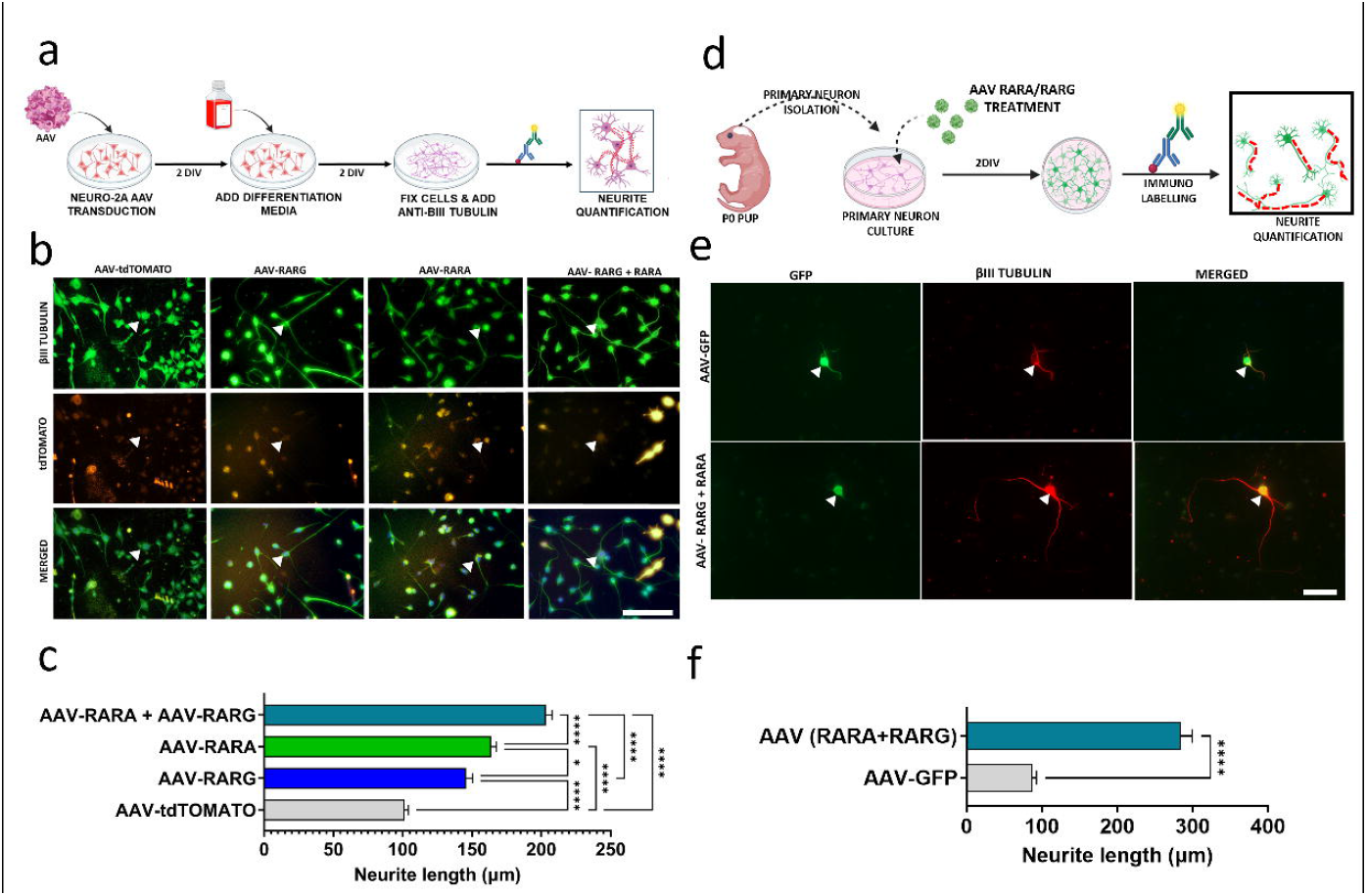
RARA and RARG synergistically promote neurite outgrowth in cultured neurons. **a.** Experimental timeline for Neuro-2A (N2A) cell culture. Cells were transduced with AAV vectors, cultured in differentiation media, fixed at 2 days *in vitro*, and stained with anti-βIII tubulin for neurite visualization and quantification. **b.** Representative immunofluorescence images of N2A cells showing GFP (viral transduction), βIII tubulin (neurites), and DAPI (nuclei) staining for control (AAV-tdTOMATO) and treatment groups (AAV-RARG, AAV-RARA, AAV-RARA + AAV-RARG). White arrowheads indicate representative neurons with extended neurites. (scale bar= 50µm) **c.** Quantification of neurite length in N2A cells across all treatment groups. Data were analysed using one-way ANOVA (F(3,1369) = 109.73, p < 0.0001), with post hoc pairwise comparisons performed using Tukey’s multiple comparison test. Combined AAV-RARA plus AAV-RARG treatment produced the longest neurites (mean 203.3 ± 76.4 μm), significantly exceeding AAV-RARA alone (163.8 μm, p < 0.0001), AAV-RARG alone (145.8 μm, p < 0.0001), and control (101.3 μm, p < 0.0001). AAV-RARA and AAV-RARG individually promoted neurite extension compared to control (both p < 0.0001). Bars represent ± SEM . **d.** Experimental design for primary neuron culture. Primary neurons were isolated from mouse brain, treated with AAV RARA and RARG , cultured for 2 days in vitro, stained with anti-βIII tubulin, and quantified. **e.** Representative immunofluorescence images showing GFP and anti-βIII tubulin staining in primary neurons treated with AAV-GFP (control) or AAV (RARA+RARG). The merged image demonstrates efficient viral transduction and robust neurite outgrowth in treated neurons. **f.** Quantification of neurite length in primary neurons comparing control versus combined RARA +RARG treatment. Data were analysed using the Mann-Whitney U test (p < 0.0001 by exact test), with AAV (RARA plus RARG)-treated neurons displaying significantly longer neurites (mean 283.97 ± 122.0 μm, n = 65 neurites) compared to control (mean 86.99 ± 39.9 μm, n = 55 neurites), demonstrating a 3.3-fold enhancement in neurite extension. Bars represent ± SEM (scale bar= 50µm) .

Together, these results show that RARA and RARG are sufficient to drive neurite outgrowth cell-autonomously, and that co-expressing both receptors produces the largest effect in both Neuro-2a cells and primary cortical neurons.

### Combined RARA and RARG expression drives cross-midline axonal sprouting after pyramidotomy injuries

Having shown that RARA and RARG gene treatment promote neurite growth *in vitro,* we asked whether this activity translates into axon growth after injury *in vivo*. We unilaterally injected AAV vectors encoding RARA, RARG, or both into layer V motor cortex neurons of the corticospinal tract (CST), and one week later transected the dominant CST projection by pyramidotomy (Fig. 3a). Overexpression of RARA and RARG viral constructs were confirmed *in-vivo* using IHC (Supplementary figure 3). We confirmed lesion completeness in every animal by the absence of anti-PKC-gamma staining in the transected pyramid (Fig. 3b), so that any labeled fibers on the denervated side of the cervical cord at 60 days post-injury reflected sprouting from the spared, AAV-transduced tract rather than incomplete transection. We quantified sprouting into the denervated cervical cord as a fiber index at three distances from the midline (200, 400, and 600 µm; Fig. 3c, d). Sprouting differed across the four groups at all three distances (Kruskal-Wallis: 200 µm, p < 0.0001; 400 µm, p < 0.0001; 600 µm, p = 0.0006). By Dunn’s post hoc test with Benjamini-Krieger-Yekutieli correction, the combined AAV-RARA + AAV-RARG group was the only one to exceed AAV-GFP control at every distance (200 µm, p = 0.0007; 400 µm, p = 0.0008; 600 µm, p = 0.0020), with fiber index means of 0.43, 0.17, and 0.08 at 200, 400, and 600 µm, respectively, versus 0.03, <0.01, and near zero in GFP controls. Neither single factor reached significance at any distance (AAV-RARA: p = 0.55, 0.37, 0.71; AAV-RARG: p = 0.13, 0.18, 0.43 at 200, 400, and 600 µm), though both showed a trend above GFP controls (n = 4 per group, except AAV-RARA + AAV-RARG, n = 5) (Supplementary table S4).

**Figure 3:**
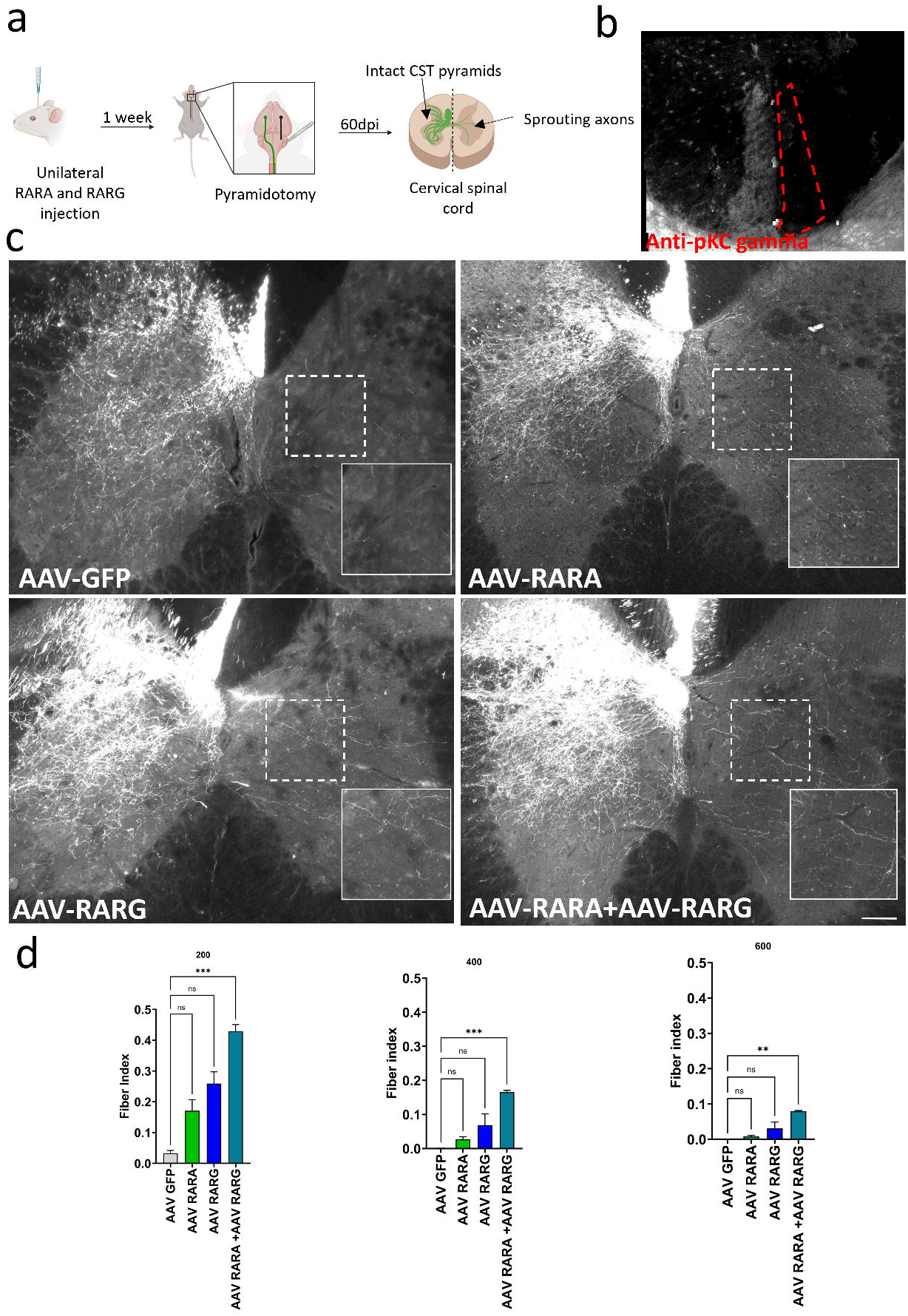
Combinatorial gene expression of RARA and RARG enhances axonal sprouting in the spinal cord. **a.** Experimental timeline showing unilateral injection of AAV-RARA, AAV-RARG and AAV-RARA+AAV-RARG into the layer V motor cortex Cortico Spinal Tract(CST) neuron, followed by pyramidotomy at 1 week, with analysis of sprouting axons in the cervical spinal cord. **b.** Image shows coronal section of spinal cord labelled with pKC-gamma , the red dotted area shows absence of pKC-gamma staining confirming completeness of pyramid surgery. **c.** Representative images of sprouting axons from the four experimental groups: AAV-GFP (control), AAV-RARA, AAV-RARG, and AAV-RARA+AAV-RARG. Dashed boxes indicate the region of interest with enlarged insets showing detail of axonal sprouting patterns. **d.** Quantification of axonal sprouting across the three measured parameters for each group. Data were analyzed using the Kruskal-Wallis test (p < 0.0001), with post hoc pairwise comparisons performed using Dunn’s test with Benjamini-Krieger-Yekutieli correction for multiple comparisons. AAV-RARA + AAV-RARG treatment resulted in significantly increased sprouting compared to control (AAV-GFP, *** p = 0.0008), while AAV-RARA alone (p = 0.37) and AAV-RARG alone (p = 0.18) showed no significant differences from control. Bars represent ± SEM with n = 4 animals per group except AAV-RARA + AAV-RARG (n = 5),(scale bar= 100µm).

Combined RARA and RARG expression therefore produces axonal sprouting after CST injury that neither receptor achieves alone, and this effect holds across the full distance range examined, extending the *in vitro* findings to an *in vivo* injury model.

### Combined RARA and RARG expression produces a detectable increase in axon regeneration after thoracic crush injury

Having shown that combined RARA and RARG expression drives sprouting after pyramidotomy, we asked whether the effect extends to axon regeneration after a more severe injury. We delivered AAV vectors expressing RARA, RARG, or both bilaterally by stereotactic injection, and one week later challenged animals with a thoracic crush injury at T8 (Fig. 4a). Overexpression of RARA and RARG viral constructs were confirmed *in-vivo* using IHC (Supplementary figure 3). Tissue was processed and imaged 12 weeks post-injury. Anti-GFAP immunostaining confirmed glial scar formation at the lesion site in all animals, indicating a consistent injury across groups (Fig. 4b) (Supplementary figure 4. At the lesion epicentre, the combined AAV-RARA + AAV-RARG condition showed denser axon labelling within the region of interest than AAV-GFP, AAV-RARA, or AAV-RARG alone (Fig. 4c).

**Figure 4:**
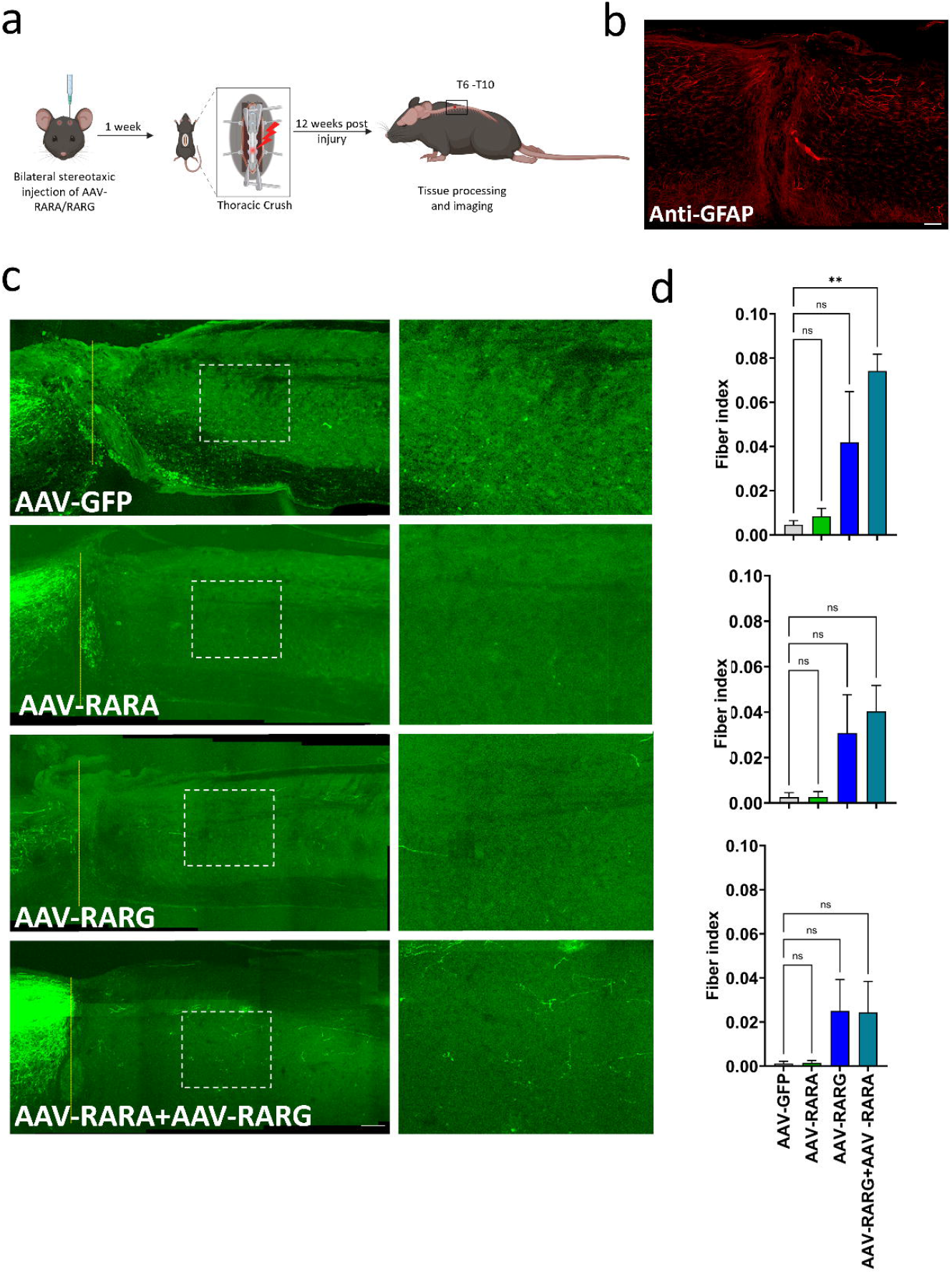
Combined RARA and RARG expression promotes axonal regeneration following spinal cord injury. **a.** Experimental design showing delivery of AAV vectors expressing RARA, RARG or both, followed by thoracic crush injury. **b.** Representative image showing anti-GFAP immunofluorescence image showing glial scar formation and confirming spinal cord injury (scale bar= 100µm). **c.** Representative images of regenerating axons at the lesion epicentre for each treatment group. Panels show spinal cord cross-sections from AAV-GFP (control), AAV-RARA, AAV-RARG, and AAV-RARA+AAV-RARG treated animals. Dashed boxes indicate the region of interest with enlarged insets displaying axonal regeneration patterns. **d.** Graph showing fiber index at 500µm, 1000µm and 1500µm from site of injury. Data were analyzed using one-way ANOVA (F(3,8) = 7.065, p = 0.0123), with post hoc pairwise comparisons performed using Dunnett’s test comparing each treatment to control (AAV-GFP) with Benjamini-Krieger-Yekutieli correction for multiple comparisons. Combined AAV-RARA+AAV-RARG treatment significantly increased regeneration compared to control (** p = 0.0098), while AAV-RARA alone (p = 0.99) and AAV-RARG alone (p = 0.15) showed no significant difference from control. Bars represent ± SEM ; n = 3 to 6 animals per group. (scale bar= 250µm).

We measured fiber index at 500, 1000, and 1500 µm from the injury site (Fig. 4d). At 500 µm, the distance closest to the lesion, one-way ANOVA showed a significant treatment effect (F(3,8) = 7.065, p = 0.0123). Mean fiber index was 0.0046 in AAV-GFP controls, 0.0084 with AAV-RARA, 0.0419 with AAV-RARG, and 0.0742 with the combination. By Dunnett’s post hoc test, only the combined group differed significantly from control (p = 0.0098); neither AAV-RARA (p = 0.99) nor AAV-RARG (p = 0.15) reached significance alone. At 1000 µm the treatment effect was not significant (F(3,8) = 3.573, p = 0.0665), although the group means followed the same order (GFP 0.0026, RARA 0.0026, RARG 0.0308, combined 0.0404; combined vs. control p = 0.0759). At 1500 µm, values were low across all groups (GFP 0.0012, RARA 0.0014, RARG 0.0250, combined 0.0243) and did not differ by treatment (F(3,8) = 1.810, p = 0.2232) (Supplementary table S5).

Combined RARA and RARG expression therefore produces a small but significant increase in axon regeneration close to the lesion after thoracic crush, with no significant effect farther out. The signal is confined to the region nearest the injury, in line with the difficulty of achieving long-distance regeneration after a severe crush. As in the pyramidotomy model, neither receptor was sufficient alone.

### Combinatorial RARA/RARG expression redirects the injured cortical transcriptome toward combination-specific targets

The *in vitro* and *in vivo* growth assays share a striking feature: neither RARA nor RARG promote significant growth individually, yet combinatorial treatment drives significant growth *in vitro* and *in vivo*. At least two regulatory models could explain this phenotype. In the first, the two receptors regulate largely the same genes, and co-delivery increases co-occupancy and transcriptional output at that shared set, so the combination is a stronger version of either factor acting on the same targets. In the second, co-delivery changes which target genes the receptors regulate, so neurons receiving both factors engage a set of targets that neither factor reaches on its own, and these combination-specific targets are the ones required for growth. The two models make opposite predictions for the transcriptome: Model 1 predicts amplified regulation of a shared gene set, whereas Model 2 predicts a distinct, combination-specific set of DEGs.

To distinguish the two models, we profiled the injured motor cortex by single-nucleus RNA sequencing (snRNA-seq), using a pooled, multiplexed design that assigns groups within the same tissue and removes batch and animal-to-animal variation. Each animal (n = 5-6) was co-injected with AAV-GFP, AAV-RARA, and AAV-RARG together, underwent pyramidotomy one week later, and the injected motor cortex was isolated one week post-injury and pooled (Fig. 5a). Nuclei were FACS-sorted on GFP signal to collect the transduced population, and each GFP+ nucleus was then assigned to a group by its transgene expression: GFP alone (control), GFP with RARA, GFP with RARG, or GFP with both RARA and RARG (combination). Because all four groups derive from the same pooled, GFP-sorted tissue, differences between them reflect transgene content rather than batch or surgical variation. UMAP feature plots confirmed the expected transgene expression in each group (Fig. 5b) (Supplementary figure 5).

**Figure 5.**
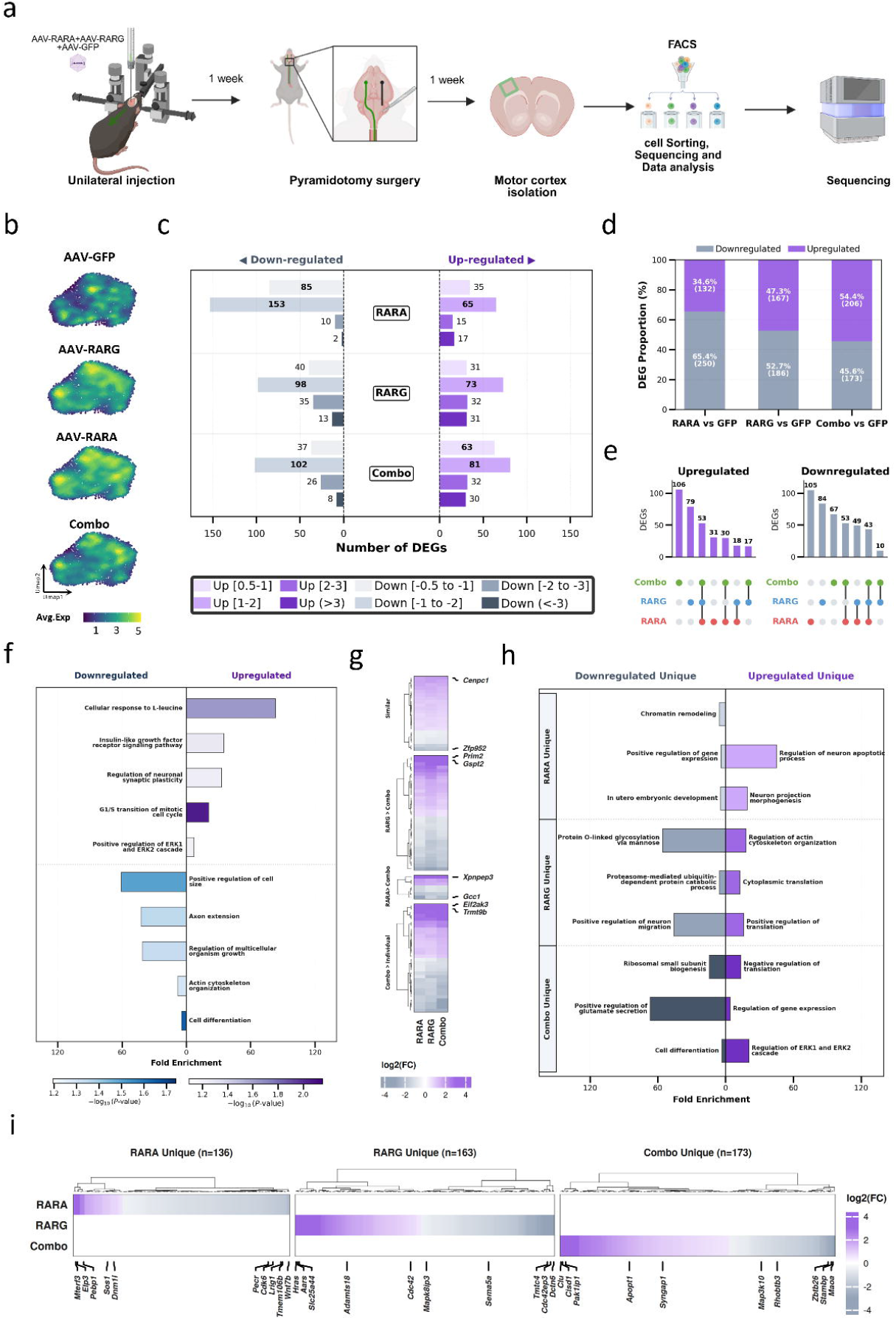
Single-nucleus transcriptomic profiling of the mouse motor cortex following AAV-mediated overexpression of RARA, RARG, and combined treatments. Across all panels, upregulated genes and associated biological processes are denoted by shades of purple, while downregulated elements are denoted by shades of grey. **a.** Schematic overview of the experimental design. Mice received unilateral injections of a combination of AAV-GFP, AAV-RARA and AAV-RARG. Following a one-week incubation, animals underwent pyramidotomy surgery. Motor cortices were isolated one week post-injury, followed by fluorescence-activated cell sorting (FACS) and single-nucleus RNA sequencing (snRNA-seq). **b.** UMAP feature plots displaying the average expression (Avg.Exp) of the respective transgenes across the four experimental groups. **c.** Bar chart summarizing the total number of upregulated and downregulated differentially expressed genes (DEGs) for the RARA, RARG, and Combo groups relative to the GFP control. Significance thresholds were defined as p.value 0.05 with a log_2(Fold Change) (≥) 0.5 for upregulation and (≤) -0.5 for downregulation. Color intensity corresponds to the magnitude of the fold change. **d.** Stacked bar graphs illustrating the overall proportion (%) of upregulated versus downregulated DEGs within each experimental condition. **e.** UpSet plots detailing the intersection of shared and unique upregulated (left) and downregulated (right) DEGs across the three overexpression groups. **f.** Gene Ontology (GO) enrichment analysis (performed via DAVID) of the DEGs common to all three treatment groups. Bar length indicates fold enrichment. **g.** Heatmap visualizing the log_2(FC) of DEGs categorized by distinct regulatory patterns: expression changes that are similar across groups, RARG > Combo, RARA > Combo, and Combo > individual treatments. The color scale indicates the degree of upregulation (purple) or downregulation (grey). **h.** GO enrichment analysis highlighting the biological processes associated with the uniquely upregulated and downregulated genes in the Combo, RARG, and RARA groups. **i.** Heatmap showing the hierarchical clustering and expression profiles (log_2(FC)) of unique DEGs identified in the RARA (n=136), RARG (n=163), and Combo (n=173) groups.

Relative to the GFP-only control, RARA, RARG, and the combination each produced hundreds of differentially expressed genes (DEGs; |log2FC| ≥ 0.5, p < 0.05; Fig. 5c). The direction of regulation shifted with transgene content (Fig. 5d) (Supplementary table S6): RARA was predominantly repressive, with 65.4% of its DEGs downregulated (250 down, 132 up); RARG was more balanced (52.7% down; 186 down, 167 up); and the combination was the only group with more upregulated than downregulated genes (54.4% up; 173 down, 206 up). If the combination simply amplified a shared program, its DEGs should overlap heavily with the single-factor sets and preserve their direction. Instead, the combination reversed the overall balance toward activation, the first indication that co-delivery changes the transcriptional program rather than scaling it.

The overlap structure supported this directly. Most DEGs were unique to one group rather than shared across all three (UpSet analysis, Fig. 5e) (Supplementary table S6), and the largest single-group sets belonged to the combination. When we grouped DEGs by their pattern across groups (Fig. 5g), a distinct class of genes was regulated more strongly by the combination than by either factor alone, alongside separate RARA-dominant and RARG-dominant classes. Hierarchical clustering resolved the unique DEGs into non-overlapping RARA-unique (n = 136), RARG-unique (n = 163), and combination-unique (n = 173) sets, each with its own expression profile (Fig. 5i).

A shared core program was still present, but it was not the growth-associated one. Among DEGs common to all three groups (Fig. 5f) (Supplementary table S7), upregulated genes were enriched for cellular response to L-leucine, insulin-like growth factor receptor signaling, regulation of neuronal synaptic plasticity, the G1/S transition, and positive regulation of the ERK1/ERK2 cascade, while downregulated genes were enriched for positive regulation of cell size, axon extension, regulation of multicellular organism growth, actin cytoskeleton organization, and cell differentiation. This common program downregulated canonical axon-growth and cytoskeletal terms and was shared by the two single-factor conditions that produced little growth on their own, so the regulation that tracks with the growth phenotype lies not in this shared set but in the combination-specific one.

The unique genes in each group carried distinct functional signatures (Fig. 5h) (Supplementary table S7). RARA-unique genes were enriched for regulation of neuron apoptotic process and neuron projection morphogenesis among upregulated terms, and for chromatin remodeling and positive regulation of gene expression among downregulated terms. RARG-unique genes were enriched for regulation of actin cytoskeleton organization, cytoplasmic translation, and positive regulation of translation among upregulated terms, and for protein O-linked glycosylation and positive regulation of neuron migration among downregulated terms. The combination-unique genes were enriched for regulation of the ERK1/ERK2 cascade and regulation of gene expression among upregulated terms, and for ribosomal small-subunit biogenesis, positive regulation of glutamate secretion, and cell differentiation among downregulated terms. Their upregulated set also included negative regulation of translation, consistent with the accompanying downregulation of ribosomal small-subunit biogenesis, indicating that the combination-specific program engages translational as well as transcriptional regulators.

Taken together, the transcriptomic data favour the second model. Co-delivery of RARA and RARG does not mainly amplify a shared set of targets. It generates a combination-specific set of differentially expressed genes, with ERK1/ERK2 signaling and gene-expression control as its defining upregulated features, that neither factor produces alone. This offers a transcriptional explanation for why the individual factors have little effect in the growth assays while the combination is active: the growth-associated regulation appears only when both receptors are present, whereas the program shared across all conditions, including the ineffective single-factor ones, downregulates canonical growth terms. Whether this reflects a change in receptor binding, as the second model predicts, is a question about occupancy rather than expression. We address it next by mapping RARA and RARG binding genome-wide.

### RARA and RARG occupy distinct genomic compartments and preferentially bind combination-specific target genes

The transcriptomic data showed that co-delivery of RARA and RARG generates a combination-specific set of target genes, and predicted that this reflects a difference in where the two receptors act on the genome. To test this, we mapped RARA and RARG binding genome-wide by CUT&RUN, profiling each receptor individually (Fig. 6a). RARA and RARG bound largely different genomic compartments (Fig. 6b) (Supplementary table S8). More than half of RARG peaks fell at promoters (55.6%), with the rest split across distal intergenic regions (21.3%), introns (15.2%), and exons (6.7%). RARA showed the opposite distribution, with most peaks at distal intergenic (49.4%) and intronic (25.3%) regions and only 8.0% at promoters. The two receptors therefore partition the regulatory genome, RARG binding predominantly promoter-proximal regions and RARA predominantly distal regulatory elements. Metagene analysis around transcription start sites (TSS) reflected this, with RARG signal peaking sharply at the TSS and RARA signal lower and more broadly distributed (Fig. 6c) (Supplementary table S8).

**Figure 6:**
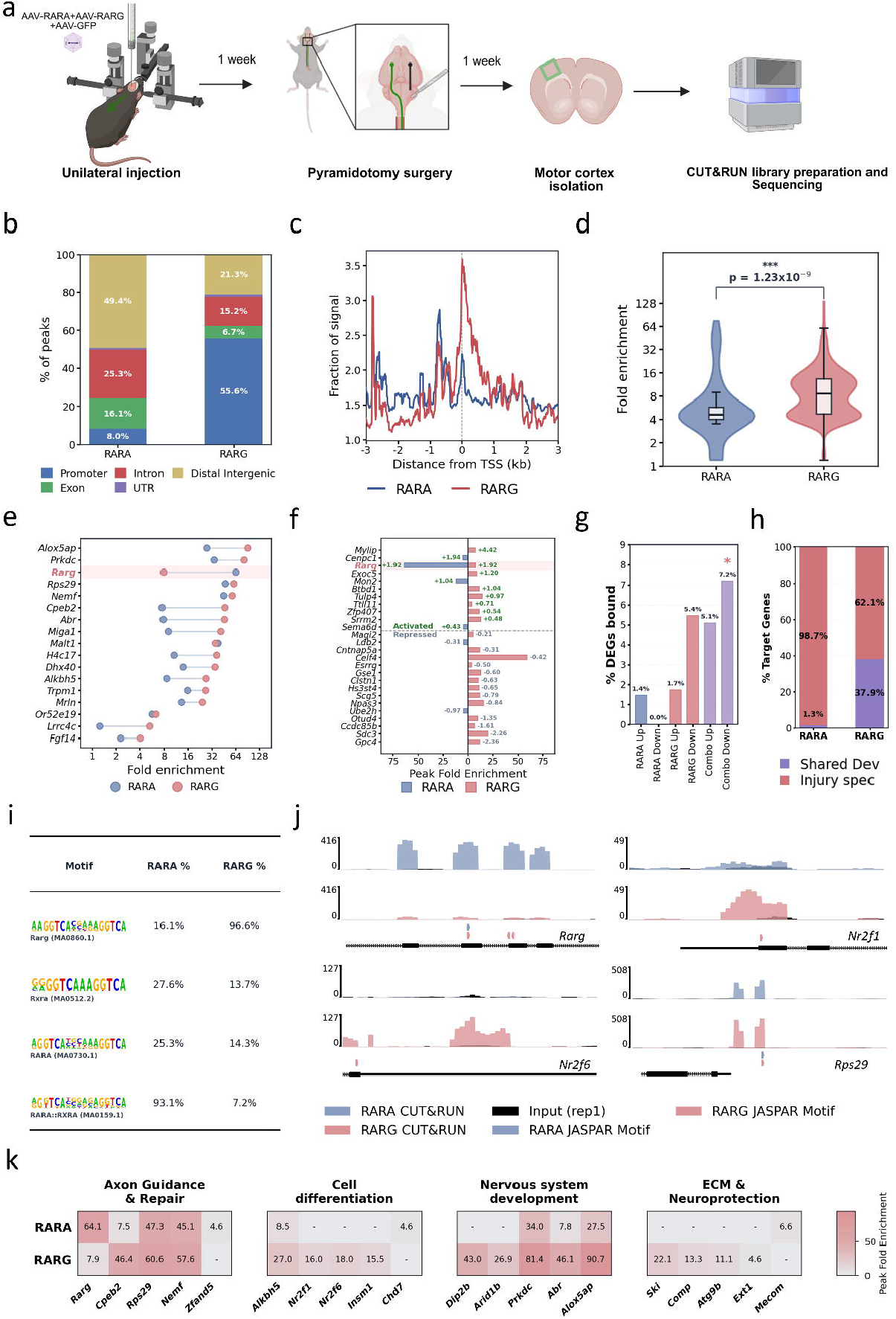
Genomic Profiling, Motif Occupancy, and Functional Characterization of RARA and RARG Binding Sites. **a.** Schematic representation of the experimental workflow: Workflow outline of CUT&RUN profiling for Retinoic Acid Receptor Alpha (RARA) and Gamma (RARG) in murine layer 5 motor cortex after pyramidotomy injury. b. Genomic distribution of RARA and RARG peaks: Stacked percentage bar plot showing the genomic distribution of RARA and RARG binding sites across promoters, introns, distal intergenic regions, exons, and UTR sites. c. TSS signal enrichment profile: Metagene line plot showing the fraction of CUT&RUN signal relative to the distance from the Transcription Start Sites (TSS) for RARA (blue line) and RARG (red line). d. Peak fold enrichment comparison: Box-and-violin plots comparing peak fold enrichment values between RARA and RARG CUT&RUN peaks. Statistical significance was determined using a two-sided Mann-Whitney U test (p = 1.23 x 10^-9^, ***). e. Differential enrichment at co-bound loci: Fold enrichment scores of combined peak regions, highlighting that RARG shows significantly higher binding enrichment at RARA-bound sites (highlighted). f. Integration of peak enrichment and transcriptomic expression: Bar plots showing MACS2 peak fold enrichment values aligned alongside corresponding single-nucleus RNA-seq (snRNA-seq) expression profiles for representative activated and repressed target genes. g. Target gene occupancy and transcriptional activation: Percentage bar plots showing direct binding rates of RARA, RARG, and co-bound (combo) targets compared against transcriptional changes in snRNA-seq datasets. h. Proportions of developmental versus adult injury-specific target genes for RARA and RARG. Stacked bars display the percentage of target genes shared with embryonic development (purple) versus adult injury-specific binding (coral) for RARA (1.3% shared, 98.7% injury-specific) and RARG (37.9% shared, 62.1% injury-specific). i. Primary Retinoid Motif Occupancy Table: Percentage of RARA and RARG peaks containing the specified monomeric or heterodimeric JASPAR motifs (Rarg MA0860.1, Rxra MA0512.2, RARA MA0730.1, RARA::RXRA MA0159.1) scanned at a 75% relative score threshold. j. Genomic locus snapshots of representative binding sites: Representative genomic track snapshots of zoomed-in peak summits (2.5 Kb windows) showing Treated samples (RARA: slate blue, RARG: rose pink; alpha 0.8) overlayed on top of their respective Input controls (solid black; alpha 1.0) sharing the same y-scale. Gene models (UCSC-style) and exact JASPAR motif match locations (80% threshold) are shown below the quantitative tracks. k. GO-Grouped Target Peak Enrichment Heatmap: Horizontal heatmap showing peak fold enrichment values for key representative genes, split and grouped across 4 major enriched Gene Ontology (GO) Biological Process pathways (Hindlimb morphogenesis, Cell differentiation, Nervous system development, and Bone morphogenesis). Gene names are in mouse standard format (title-cased and italicized, e.g. Rarg), and non-enriched cells are colored in a distinct soft gray (-).

RARG also bound more strongly. Across all peaks, RARG fold enrichment exceeded that of RARA (Mann-Whitney U test, p = 1.23×10⁻⁹; Fig. 6d). At loci bound by both receptors, RARG again showed significantly higher enrichment than RARA at the same sites (Fig. 6e) (Supplementary table S8), so even where the two co-occupy, RARG is the dominant binder. Motif analysis supported direct, sequence-specific binding by each receptor through its own motif (Fig. 6i) (Supplementary table S8): 96.6% of RARG peaks contained the RARG motif (JASPAR MA0860.1) versus 16.1% of RARA peaks, whereas 93.1% of RARA peaks contained the RARA::RXRA heterodimer motif (MA0159.1) versus 7.2% of RARG peaks. Each receptor thus binds its own preferred motif, consistent with genuine occupancy rather than shared or indirect recruitment.

To connect binding to transcriptional output, we asked what fraction of the snRNA-seq DEGs in each group were directly bound (Fig. 6g). Direct binding rates were low for the single-factor DEGs (RARA-upregulated 1.4%, RARA-downregulated 0.0%; RARG-upregulated 1.7%, RARG-downregulated 5.4%) and higher for the combination DEGs, which showed the greatest direct occupancy of any group (combination-upregulated 5.1%, combination-downregulated 7.2%). The combination-specific target genes were therefore the most likely to be direct binding targets, linking the combination-specific transcriptional program to direct genomic occupancy. Aligning peak fold enrichment with the direction of expression change showed that directly bound genes included both activated and repressed targets (Fig. 6f) (Supplementary table S8), matching the mixed activating and repressive roles of the receptors seen in the snRNA-seq data (Fig. 5f). We next compared injury binding with the embryonic binding profiles to ask whether each receptor returns to its developmental sites (Fig. 6h). The two receptors differed sharply. Only 1.3% of RARA target genes were shared with developmental sites, with the remaining 98.7% specific to the injured adult cortex. RARG was split between the two, with 37.9% of targets shared with development and 62.1% injury-specific. RARG therefore re-occupies a substantial part of the program it used embryonically, whereas RARA binds almost entirely new loci after injury.

Representative loci illustrated these patterns (Fig. 6j) (Supplementary table S8). At Rarg, Nr2f1, Nr2f6, and Rps29, RARA and RARG peaks overlaid their respective JASPAR motif positions, with the stronger RARG signal at promoter-proximal summits matching the genome-wide distribution. Grouping bound target genes by GO biological process (Fig. 6k) (Supplementary table S9) placed them in pathways relevant to the growth phenotype, including nervous system development (for example Prkdc, Abr, Dip2b, Arid1b), cell differentiation, hindlimb morphogenesis, and bone morphogenesis, with individual genes showing receptor-preferential enrichment (strong RARG binding at Prkdc and Abr, strong RARA binding at Rarg).

Together, the CUT&RUN data show that RARA and RARG occupy distinct and largely non-overlapping regions of the genome, promoter-proximal for RARG and distal for RARA, so that the two receptors cover regulatory space that neither reaches alone. Because binding was profiled for each receptor individually, we cannot directly measure how co-delivery alters occupancy; combinatorial binding can only be inferred from the overlap and complementarity of the two single-factor maps. Within that limit, the data are consistent with the transcriptomic model: the combination-specific genes were the most directly bound, and the two receptors engage complementary genomic compartments, supporting cooperation through complementary target engagement rather than amplification of a shared set.

### RARG independently promotes neurite outgrowth in peripheral sensory neurons

Having established that combined RARA and RARG expression drives sprouting and regeneration in the central nervous system, we asked whether this combinatorial requirement is a general feature of these two receptors or one that is specific to the CNS context. To test this, dorsal root ganglia (DRG) were isolated seven days after intrathecal AAV injection, cultured in 96-well plates, and neurite outgrowth was imaged and quantified at 2 days *in vitro* (Fig. 7a, b). Contrary to the pattern observed in CNS neurons, AAV-RARG alone was sufficient to significantly increase neurite length compared to AAV-GFP control (mean 301.4 µm vs. 165.6 µm; Kruskal-Wallis test, p = 0.0013, with Dunn’s post hoc test and Benjamini-Krieger-Yekutieli correction, p = 0.0007). AAV-RARA alone had no effect on neurite length relative to control (157.9 µm, p > 0.9999), and, notably, the combined AAV-RARA + AAV-RARG treatment did not reach significance over control (219.7 µm, p = 0.3859) (Supplementary table S10), despite trending higher than AAV-RARA alone or AAV-GFP. This indicates that in peripheral sensory neurons, RARG acts as an independent driver of neurite outgrowth, and that co-expression with RARA does not enhance, and may even attenuate, this single-factor effect.

**Figure 7:**
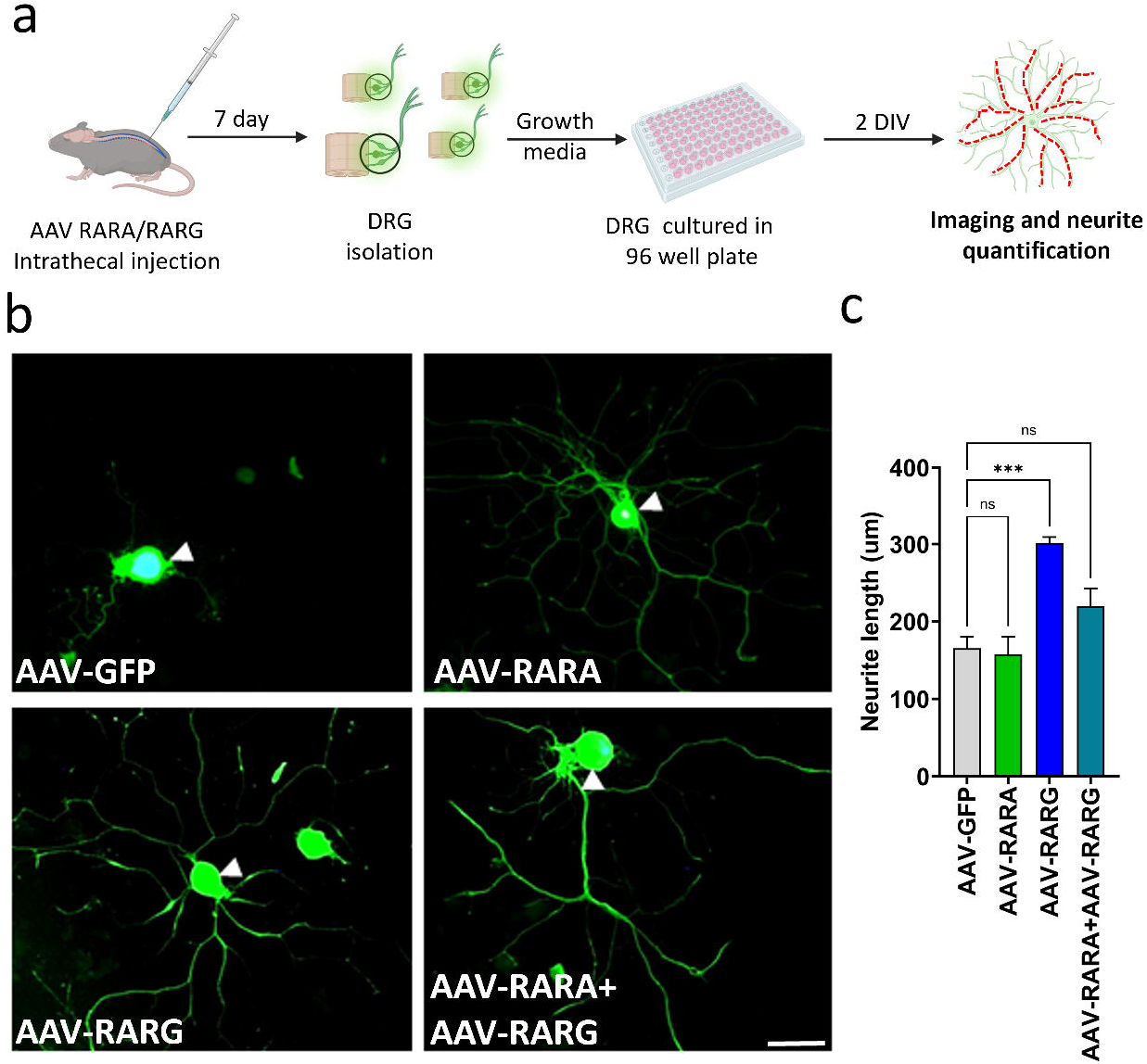
RARG promotes neurite outgrowth in peripheral sensory neurons but requires RARA cooperation for central nervous system axonal regeneration. **a.** Experimental timeline showing AAV RARA/RARG intrathecal injection into mice, followed by dorsal root ganglion (DRG) isolation 7 days later, culturing of DRG neurons in growth media within a 96-well plate, and imaging and neurite quantification at 2 days in vitro (DIV). **b.** Representative immunofluorescence images of DRG neurons expressing AAV-GFP (control), AAV-RARA, AAV-RARG, or AAV-RARA+AAV-RARG. Arrowheads indicate the neuronal soma. **c.** Quantification of neurite length (µm) for each treatment group. Data were analyzed using the Kruskal-Wallis test (p = 0.00135), with post hoc pairwise comparisons performed using Dunn’s test with Benjamini-Krieger-Yekutieli correction for multiple comparisons. AAV-RARG treatment significantly increased neurite length compared to control (*** p = 0.00074), while AAV-RARA alone (p = 1.0, ns) and AAV-RARA + AAV-RARG combination (p = 0.386, ns) showed no significant enhancement. Bars represent ± SEM ; n = 4 to 12 neurons per group. (scale bar= 100µm)

Taken together, these findings reveal that the combinatorial requirement for RARA and RARG observed in CNS neurons is not a universal feature of these receptors’ growth-promoting activity but rather a context-dependent one. While RARG appears capable of driving neurite outgrowth on its own in the peripheral nervous system, its ability to do so in the CNS instead depends on RARA cooperation, suggesting that the two receptors engage distinct or complementary mechanisms depending on neuronal identity, and that RARA’s role in the CNS may be to enable or potentiate a growth program that RARG can otherwise execute independently in a more permissive PNS environment.

### Combined RARA and RARG treatment improves locomotor recovery after thoracic crush injury

To test whether the anatomical regeneration seen with combined treatment translates into functional recovery, we assessed hindlimb locomotion after thoracic crush injury. Mice injected with AAV-RARA, AAV-RARG, AAV-RARA+AAV-RARG, or AAV-GFP control walked freely on a horizontal ladder, and their movements were recorded at 30 fps weekly from 1 to 12 weeks post-injury (WPI) (Fig. 8a). We extracted sub-pixel coordinates for six hindlimb markers using DeepLabCut, generated stance–swing stick plots with a custom script, and derived spatiotemporal gait parameters for longitudinal analysis.

**Figure 8:**
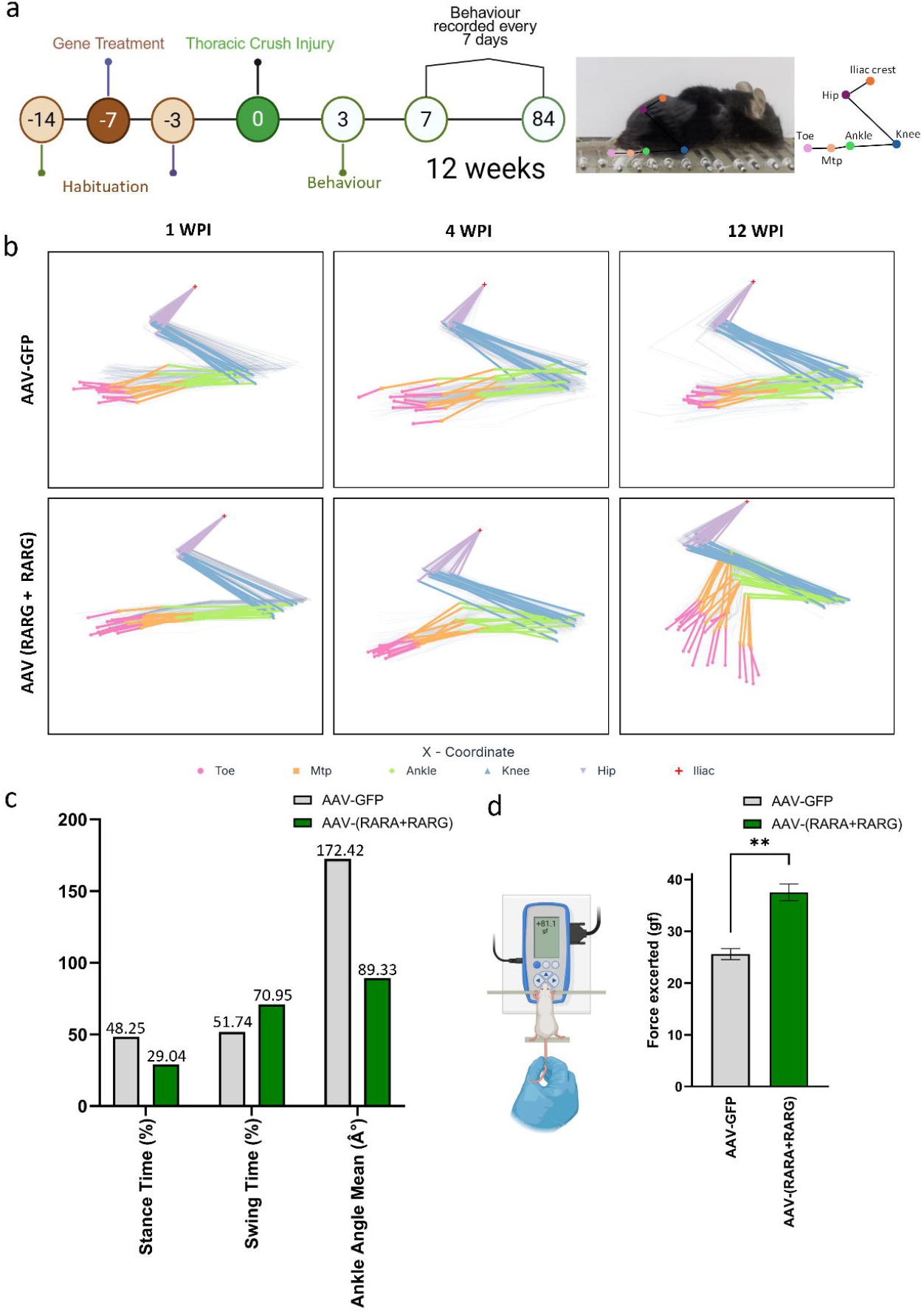
Behavioural and kinematic analysis following thoracic crush injury and gene therapy. **a.** Schematic workflow showing experimental design for habituation, anterograde injection of AAV-GFP and AAV-RARA+AAV-RARG followed by thoracic crush injury and gait analysis to assess functional recovery. **b.** Hind limb gait analysis plots of 1 week post injury and 12 week post injury of the treatment and control groups to visualise the extent of recovery over a period of 12 weeks. **c.** Graph comparing the kinematic parameters (stance time (%), swing time (%) and ankle angle mean (Â°)) between control and AAV-RARA+AAV-RARG treated groups at 12 weeks post injury time point. **d.** Illustration of the grip strength meter setup used to assess the hind limb grip strength at 12 weeks post injury time point followed by a comparison graph quantifying the hindlimb force exerted (gf) by control and AAV-RARA+AAV-RARG treated groups (Welch’s t-test, p = 0.0055; Bars represent ± SEM ; n = 3 per group).

Stick-plot trajectories showed a progressive restoration of the hindlimb stepping envelope in the combination-treated group (Fig. 8b) (Supplementary table S11). Uninjured locomotion follows a three-part stride: a posterior backstroke, a vertical lift to clear the rungs, and an anterior forward swing. AAV-GFP control animals showed a severe, persistent deficit across the 1–12 WPI period, with flattened trajectories consistent with continuous hindlimb dragging. By 12 WPI, AAV-RARA+AAV-RARG animals re-established the early components of the gait cycle, executing a coordinated backstroke and a pronounced vertical lift that eliminated rung dragging; the anterior swing remained partially incomplete, with limbs often elevated and replaced near their point of origin rather than translating fully forward, but a substantial portion of the normal stepping trajectory was recovered.

Because the combination was the only group to show clear trajectory recovery, we restricted quantitative gait profiling to AAV-RARA+AAV-RARG versus AAV-GFP controls at 12 WPI (Fig. 8c) (Supplementary table S11). The combination group showed a shift toward a normalized weight-bearing pattern, with stance time reduced from 48.25% to 29.04% and swing time increased from 51.74% to 70.95%. Mean ankle angle was also lower in the combination group (89.33°) than in controls (172.42°), indicating restored joint flexion rather than the rigid, extended posture seen after injury.

Functional strength assessment agreed with the gait findings (Fig. 8d) (Supplementary table S11). Using a grip-strength meter, AAV-RARA+AAV-RARG animals exerted greater hindlimb grip force at 12 WPI than AAV-GFP controls (Welch’s t-test, p = 0.0055; n = 3 per group). Combined RARA and RARG treatment therefore improves both gait kinematics and hindlimb strength after thoracic crush injury.

## Discussion

Here we identify two retinoic acid receptors, RARA and RARG, as a cooperative pair of transcription factors that reactivate axon growth in the injured corticospinal tract, and we define the logic of their cooperation. Neither receptor was sufficient alone in any assay, from neurite outgrowth in Neuro-2a cells and primary cortical neurons to corticospinal sprouting after pyramidotomy, axon regeneration after thoracic crush, and the accompanying recovery of gait and grip strength. Single-nucleus transcriptomics and CUT&RUN indicate why. The two receptors regulate a combination-specific set of genes rather than a shared set, occupy complementary genomic compartments, and, in the case of RARG, show a binding-to-expression relationship that resolves only when both are present. The sections below develop these observations and place them against the emerging role of nuclear-receptor transcription factors in CNS repair.

Combinatorial transcription factor delivery is one way to reactivate the growth programs that adult CNS neurons lose after development, but few cooperative pairs have been validated, and the mechanistic basis for their cooperation is rarely resolved ^9,12^. Our data address both points for the retinoic acid receptors. Across every growth assay, RARA and RARG were effective together but not alone. Two models could explain this. In the first, the receptors regulate a common set of genes, and co-delivery raises transcriptional output at those shared targets. In the second, co-delivery changes which genes are engaged, so the neuron activates a target set that neither receptor reaches individually. Our data favour the second model. The combination generated its own set of differentially expressed genes rather than amplifying the single-factor programs (Fig. 5e,g,i). The program common to all three conditions, including the two ineffective single-factor conditions, downregulated rather than upregulated the canonical axon-growth terms (Fig. 5f), so the regulation that tracks with growth lies in the combination-specific set. At the level of binding, RARA and RARG occupied largely distinct genomic compartments, RARG at promoters and RARA at distal regions (Fig. 6b), covering regulatory space that neither occupies alone. Because CUT&RUN was performed for each receptor individually, we cannot measure how co-delivery alters occupancy directly; the combinatorial binding state is inferred from the complementarity of the two single-factor maps. Within that limit, the transcriptomic and genomic data point to the same conclusion: RARA and RARG cooperate by engaging complementary targets, and the combination-specific targets are the ones associated with growth. Cooperation through target redistribution rather than target amplification is consistent with reports that transcription factor pairs, including KLF6 with NR5A2 and CTCF with YY1, promote axon regeneration only in combination^9,13^.

The requirement for both receptors was specific to central neurons. In dorsal root ganglion neurons, RARG alone was sufficient to drive neurite outgrowth, and adding RARA gave no further benefit (Fig. 7). This dissociation is informative for two reasons. Peripheral sensory neurons have long served as a discovery platform for factors later tested in the CNS, on the assumption that intrinsic growth programs are broadly shared^2^ ; our results show that the same two factors are wired differently in the two systems, so a mechanism defined in DRG neurons does not transfer directly to corticospinal neurons. The RARA requirement is a central feature, not a general property of RAR-driven growth. The contrast also indicates where the combinatorial mechanism operates. If RARG alone reconstitutes the growth state in peripheral neurons but requires RARA centrally, the distal, RARA-bound elements identified by CUT&RUN are candidates for the central-specific component of the program. Whether this reflects differences in chromatin accessibility, cofactor availability, or receptor stoichiometry between the two neuronal classes is not resolved by our data and will require occupancy profiling in both cell types.

One combination-specific event stands out and connects binding to expression. RARA bound the RARG locus at 370-fold enrichment (Fig. 6), and RARG was upregulated only in the combination (avg log₂FC +1.92, adjusted p = 1.1×10⁻¹⁰; Fig. 5), not by RARA or RARG alone (adjusted p = 1.0 in each). This pattern suggests that part of RARA’s activity is relayed through RARG: RARA occupies the RARG locus, but RARG transcript rises only when both receptors are present. A relay of this kind has a precedent in the canonical retinoid cascade, in which liganded RARA drives transcription of downstream receptors ^14^. The established downstream target there is RARB2 rather than RARG, and RARB was not induced in our data (adjusted p = 1.0 in all conditions), so a direct RARA-to-RARG relay appears to be a neuron-specific variant of that cascade rather than the classical one. It also offers one explanation for why RARA alone is insufficient, since its downstream effect depends on a partner it helps induce but cannot induce by itself.

Beyond RARG, several CUT&RUN targets place the receptors at genes controlling translation and ribosome function. RARG bound Celf4 (60-fold), a neuronal 3′UTR-binding protein of the CELF family whose members regulate axon regeneration through control of mRNA translation and splicing^15,16^, Nemf and the ribosomal protein gene Rps29 (each ∼50-fold), with NEMF a core component of ribosome-associated quality control whose loss causes motor neuron degeneration^17^; and Ncoa1 (20-fold), which encodes the p160 coactivator SRC-1 that RARs use to recruit histone acetyltransferases^18^. These genes were not called as differentially expressed in the snRNA-seq data, but that absence is difficult to interpret as true transcriptional silence. Single-nucleus sequencing samples nuclear RNA at shallow per-cell depth and reliably detects only abundant transcripts, so moderately expressed genes such as these are prone to false negatives. Two readings are therefore compatible with the data: the receptors bind these loci without changing their transcription, or they change it below the detection threshold of the assay. Both are consistent with an effect exerted post-transcriptionally, which is the mode of action of the bound targets themselves, since CELF proteins, NEMF, and ribosomal components act on translation rather than transcription. This interpretation aligns with the combination-specific program at the pathway level, which was enriched for negative regulation of translation and depleted for ribosomal small-subunit biogenesis (Fig. 5h). It is also notable that ribosomal protein genes are downregulated roughly 56-fold during maturation of CNS projection neurons, in step with the developmental decline in intrinsic growth capacity, and that their experimental re-upregulation promotes axon regeneration^19^. Receptor occupancy at translational-control loci, together with a combination-specific translational signature, indicates that combinatorial RAR activity engages this axis whether or not it registers as a change in bulk transcript level.

The two receptors also differed in how their injury binding related to their developmental binding. Relative to the embryonic profiles, RARA binding after injury was almost entirely new: 98.7% of RARA target genes were injury-specific and only 1.3% were shared with developmental sites (Fig. 6h). RARG was split, with 62.1% injury-specific targets and 37.9% shared with developmental loci. RARG therefore re-occupies a substantial part of the developmental program it used embryonically, whereas RARA is redeployed almost entirely to new locations after injury. This asymmetry fits the rest of the data. RARG behaves as the receptor that reuses the developmental growth program, which is consistent with its sufficiency in peripheral sensory neurons, while RARA contributes a largely injury-specific, distal binding component that is required only in central neurons. The developmental origin of much of the RARG program also connects this system to the broader observation that CNS regeneration involves partial reactivation of developmental transcriptional states^2^.

This translational signature is worth placing in a broader context. We recently found that the nuclear receptors NR2F1 and NR2F6 promote corticospinal regeneration through complementary mechanisms, with NR2F6 imposing a broad translational down-shift through a conserved corepressor domain^12^. The combination-specific program described here, enriched for negative regulation of translation and depleted for ribosomal biogenesis, points to a similar theme for RARA and RARG. Post-transcriptional and translational control is increasingly linked to CNS regenerative capacity, from CPEB1-mediated stabilization of growth transcripts to RSK-driven ribosomal control of regeneration-associated protein synthesis^19–21^. The recurrence of translational regulation across independent nuclear-receptor transcription factors raises the possibility that translational reprogramming is a shared node through which this family influences axon growth, rather than a feature unique to one receptor pair. This proposal rests on GO enrichment of binding and expression data and remains to be tested directly, for example by ribosome profiling of neurons expressing RARA and RARG together.

Overall, our data identify RARA and RARG as a cooperative pair of transcription factors that reactivate axon growth in the injured central nervous system, and define the logic of their cooperation. The two receptors are ineffective alone and effective together because they engage complementary, largely non-overlapping targets rather than amplifying a shared program, with RARA relayed in part through combination-specific induction of RARG, and with RARG reusing a developmental binding program that RARA does not. This cooperation is required in corticospinal neurons but not in peripheral sensory neurons, which places the RARA requirement as a central rather than universal feature of RAR-driven growth. Together with the convergence of this receptor pair and the NR2F receptors on translational control, these findings position the nuclear-receptor family as a source of cooperative, combinable regulators of CNS regeneration, and suggest that combinatorial delivery guided by complementary genomic binding is a route to reactivating the developmental growth program in adult neurons. The main limitations are the modest regeneration achieved after complete thoracic crush, the small *in vivo* cohorts, and the inference of combinatorial binding from single-factor maps; testing predicted co-occupancy directly, and improving the magnitude and distance of regeneration, are the next steps.

## METHODS

### Animal husbandry

All the procedures done on animals were Institutional Animal Ethics Committee (IAEC) approved at CSIR-CCMB. The experimental animals, i.e., wild-type C57BL/6J mice, were housed under a 12:12-hour light/dark cycle.

### Cloning strategy

Mouse RARA [Horizon Discovery, MMM1013-202762042] and RARG [Horizon Discovery, MMM1013-202763945] were isolated from a cDNA library of mouse transcription factors. RARA and RARG were cloned using traditional cut and paste cloning using 3’ EcoRI [NEB, #R0101S ] and 5’ XbaI [NEB, #R0145S ]. The tdTOMATO reporter gene was excised from the plasmid backbone and replaced with the amplified gene using T4 DNA ligase [Takara, #2011B]. Successful cloning was confirmed by Sanger sequencing (CCMB facility) using CAG-forward (5’-GCAACGTGCTGGTTATTGTG-3’) and Amp-reverse (5’-ATAATACCGCGCCACATAGC-3’) primers.

### AAV production

HEK293T cells were used to produce Adeno-associated viral particles (serotype 9), and the AAV purification kit [Takara, #6675] was used to purify the virus. HEK293T cells were transiently transfected with pAAV-CAG-RARA or pAAV-CAG-RARG, pAAV2/9n and pAdDeltaF6 plasmids using polyethylenimine 40k [Polysciences, #24765]. The cells were harvested 72 hours post-transfection, and the virus was isolated according to the kit’s instructions. The integrity of the AAV capsid was assessed by 10% SDS-PAGE, run for 120 minutes at 35 mA, which was then followed by staining with Coomassie Blue and destaining overnight in destaining solution (Methanol-acetic acid solution), which confirmed intact capsid formation. Quantification PCR was done to assess the viral titer of the AAV particles. To remove the residual plasmid DNA on the AAV particles, DNase I [Thermofisher, #EN0521] treatment was done at 37℃ for 30 minutes, and then the samples were subjected to a 1:10 serial dilution, repeated four times. TB green [Takara, #RR82WR] was used to estimate viral titers by comparing with a plasmid standard curve. A titer of 10^11^ viral particles/µl was achieved for all AAV preparations.

### Primary Neuron Culture and Transduction

Poly-D-Lysine [Gibco, #A3890401] coating was performed at 0.2mg/ml on autoclaved coverslips, which were acid-treated, placed in a 12-well plate, and incubated for 2 hours at 37°C. The coverslips were then washed with autoclaved Milli-Q water and then air-dried. Motor cortex was isolated from C57BL/6J pups, adhering to IAEC guidelines, and then transferred into pre-warmed HBSS (Hank’s Balanced Salt Solution), minced, and then incubated for 10 minutes at 37 °C in 0.1% Trypsin-EDTA [Sigma, #T4799]. Fetal Bovine Serum [Gibco, #10438026] was used to inactivate trypsin, followed by trituration in plating media (Neurobasal medium [Gibco, #21103049], 5% Fetal Bovine Serum, 2% B27 supplement [Gibco, #17504044], 1X PenStrep). The cells were quantified and seeded at a density of 70,000 cells per well in 1 milliliter of plating media. Twenty-four hours post-seeding, a 50% media exchange was performed using fresh medium supplemented with 1 µM Ara-C [Sigma, #C1768]. Co-transduction was performed using pAAV-CAG-GFP and the virus of interest in a 1:2 titer ratio. At 50 hours post-transduction, efficiency was determined via immunostaining for βIII-Tubulin [CST, #2128S] to verify neuronal identity and viral uptake.

### Neuro-2a culture for in vitro neurite outgrowth assays

For the *in vitro* neurite outgrowth assays, Neuro-2a cells were maintained in Dulbecco’s Modified Eagle Medium (DMEM; Gibco, #11885084) enriched with 10% FBS and 1X penicillin-streptomycin-gentamicin (PSG). We seeded 15,000 cells per well into 96-well plates. After a 24-hour incubation, the cells underwent transduction with AAV-RARA, AAV-RARG, or AAV-RARA+AAV-RARG, alongside a tracer AAV-GFP (3:1 ratio). Following a 48-hour incubation, the cells were trypsinized using 10 μL of 0.1% trypsin (Sigma, #T4799) and transferred onto 18 mm x 18 mm glass coverslips in 12-well plates. Once adhered, the medium was swapped for a differentiation medium (DMEM and PSG) for an additional 48 hours to induce neurite outgrowth. Cells were subsequently fixed with 1 mL of 4% paraformaldehyde (Sigma, #30525-89-4) for 15 minutes before undergoing immunostaining. Imaging was conducted on a Zeiss Apotome.2 microscopy system at 20x magnification using fluorescent markers: βIII-tubulin (1:500 dilution; TRITC filter: ex 557/576 nm, em 580/620 nm), GFP (FITC filter: ex 475/495 nm, em 510/525 nm), and nuclei (DAPI filter: ex 357/44 nm, em 447/60 nm). To quantify neurite length, we traced polylines along the neurites of transfected cells on overlay images, examining 150 cells for each condition (Supplementary Table S2). All data analysis was completed using GraphPad Prism (version 10.1.0), with statistical significance determined via one-way ANOVA and subsequent unpaired t-tests. The results are reported as mean ± SEM.

### Animal surgeries

All animal experimentation protocols were authorized by the Institutional Animal Ethics Committee (IAEC) at CSIR-CCMB and strictly followed established IAEC guidelines. Experiments utilized adult C57BL/6J mice of both sexes: those >12 weeks old (20–25g) were subjected to cortical injections and thoracic crush injuries, whereas 6–8-week-old mice were used for pyramidotomy. All animals were anesthetized using a mixture of Ketamine (50 mg/ml) and Xylazine (100 mg/ml).

For cortical injections, the skull was accessed via incision to target four sites relative to Bregma (M/L: ±2 mm, A/P: 0 and +1 mm). Following craniotomies at these positions, viral vectors were administered via Hamilton syringe at a depth of 0.50–0.60 mm (D/V).

Thoracic crush injuries were performed by exposing the spine through a dorsal midline incision and blunt muscle dissection. Mice were secured in a specialized spine stabilizer to perform a T6– T10 laminectomy with spring scissors. The spinal cord was then compressed with blunt forceps (1 mm width), applying uniform force.

Pyramidotomy procedures were done by performing a ventral midline incision between the jaw and forelimbs to gain access to the medullary pyramid, followed by the partial resection of the occipital bone. After incising the dura, the right pyramidal tract was severed using iridectomy scissors (0.5 mm width, 0.25 mm depth). The incision was closed by re-aligning the trachea, oesophagus, and musculature before suturing the skin.

### Axon-growth quantification

For the pyramidotomy CST sprouting quantification, we fixed the tissue using 4% paraformaldehyde (PFA) at 4°C overnight followed by embedded in 12% gelatin (Sigma Aldrich, G2500) in PBS, and 50 µm coronal sections of the spinal cord were obtained using a vibratome and imaged using the SP8 confocal microscope. The sprouting quantification was then done using DeNAT^22^ and the resulting values were normalized against medullary pyramid GFP labelled axon counts. For thoracic crush injury experiments, we processed tissue samples following the same protocol used for pyramidotomy. We imaged 50 μm sections spanning the injury site using the STED microscope. The lesion center was defined as the location of peak GFAP immunoreactivity; using this point as a baseline, we drew reference lines 500, 1,000, and 1,500 μm in the caudal direction. We then counted the GFP⁺ axons intersecting these lines and normalized the resulting values against medullary pyramid GFP labelled axon counts.

### Immunohistochemistry

Following anesthesia, adult animals were decapitated for the spinal injury experiments; the brain and spinal cord were then harvested and post-fixed overnight in 4% paraformaldehyde (PFA) at 4°C. Spinal cord tissue was embedded in 12% gelatin (Sigma Aldrich, G2500) in PBS, and 50 µm sagittal sections were obtained using a vibratome. Sections were permeabilized in PBST and blocked with 5% normal goat serum. Subsequently, samples were incubated overnight at 4°C with primary antibodies: anti-GFAP (Cell Signalling Technology, mAb #3670, 1:500) to assess injury severity and anti-PKC gamma (GTX107639) to verify pyramidotomy success in spinal cord sections, or anti-RARA and anti-RARG (#62294, #8965 respectively) for brain motor cortex samples to evaluate overexpression. After washing, sections were incubated for 2 hours at room temperature with Alexa Fluor-conjugated secondary antibodies (Thermo Fisher, 1:500). Finally, samples were counterstained with DAPI, mounted, and imaged using high-resolution STED microscopy.

### CUT&RUN assays

Seven days following pyramidotomy injury, motor cortices were collected from mice previously injected with AAV-RARA and AAV-RARG. The tissue was minced on ice and fixed for 2 minutes in 0.1% formaldehyde (CST, #12606P), followed by quenching with 10X glycine (CST, #7005S) and subsequent washing. CUT&RUN procedures were performed using the DNA– Protein Interaction Assay Kit (CST, #86652). Briefly, cells were dounced in wash buffer containing spermidine (CST, #27287) and protease inhibitor cocktail (CST, #7012), attached to Concanavalin-A magnetic beads, and incubated overnight at 4 °C with anti-RARA & anti-RARG antibodies (#62294, #8965 respectively). After washing, bead–nuclei complexes were incubated for 1 hour in digitonin buffer (CST, #16359) containing spermidine and protease inhibitors, supplemented with pAG-MNase (CST, #15338). Digestion was activated by adding CaCl₂ for 30 minutes at 4 °C, then halted using the kit’s Stop Buffer (CST, #48105) combined with RNase A (CST, #7013). Cross-links were reversed by incubation with 10% SDS and Proteinase K (CST, #10012) at 65 °C for 2 hours, and DNA was isolated using kit spin columns (CST, #14209). Input controls were processed in parallel by lysing nuclei in DNA Extraction Buffer (CST, #42015) and sonicating on a Covaris M220 (peak power 75, duty cycle 10%, 200 cycles per burst, 75 seconds) to generate fragments of 100–600 bp. Libraries for both CUT&RUN and input samples were constructed with the NEBNext® Ultra™ II DNA Library Prep Kit (NEB, #E7103) and sequenced to achieve approximately 5 million paired-end reads per sample.

### Single-nucleus RNA-seq processing

Motor cortices were harvested from C57BL/6J mice (6–8 weeks old, mixed sex) seven days after pyramidotomy injury. One week prior to the injury, animals received anterograde injections of a combination of AAV-GFP, AAV-RARA and AAV-RARG (0.6 µL per site) at four stereotaxic coordinates relative to Bregma (M/L: ±2 mm, A/P: 0 and +1 mm). Isolated tissue was immediately flash-frozen and maintained at –80 °C. For nuclei isolation, tissue was dounced in 2 mL Nuclei Lysis Buffer supplemented with RNase inhibitor. The resulting suspension was centrifuged (500 g, 2 minutes, 4 °C), and the pellet was resuspended in Nuclear Suspension Buffer (PBS, 0.1% BSA, RNase inhibitor) before filtration through a 20 µm mesh. GFP⁺ nuclei were isolated via flow sorting on a BD Melody system (80 µm nozzle), utilizing FSC/SSC gating to exclude debris and doublets, and collected into resuspension buffer (approximately 10000 nuclei per sample). Single-nucleus libraries were prepared using the Chromium Next GEM Single Cell 3’ Kit v3.1 [10x Genomics, #PN1000121] in accordance with the manufacturer’s instructions.

### Behavioral assessments

To evaluate motor function, mice underwent training on a horizontal ladder-walking task. Animals were familiarized with the apparatus in two sessions, occurring 7 days prior to and 3 days after stereotaxic AAV administration, to mitigate any effects of environmental novelty. The ladder consisted of an aluminium frame (23 × 2 × 6 inches) with rungs spaced at 1-cm intervals, flanked by clear acrylic walls to maintain the subject’s position within the field of view. Starting one week post-injury and continuing weekly until the twelfth week, each mouse completed three consecutive traversals per session. The videos were recorded at 30 fps using a side-mounted camera aligned with the ladder’s midline.

For initial data processing, AVI files were analyzed using ImageJ; every tenth frame was evaluated by measuring the vertical distance between the greater trochanter and a baseline connecting the tarsal joints. This calculated “mean hip-lift” value allowed for the differentiation between active limb movement and passive dragging. For more granular kinematic analysis, videos were processed via DeepLabCut v3.1. We manually annotated fifty frames per dataset to define six anatomical landmarks: iliac crest, hip, knee, ankle, MTP joint, and toe. A ResNet-50 network was trained until the mean pixel error was reduced to below 2 px. The resulting model provided time-resolved x–y coordinates for each landmark, which were subsequently analyzed using Python scripts to quantify stance and swing phases, toe-clearance trajectories, and inter-joint angles.

### Intrathecal delivery of RARs

For the delivery of viral vectors via intrathecal injection, adult C57BL/6J mice(mixed sex, >6 weeks old, 20–25 g) were anesthetized using a mixture of Ketamine (50 mg/ml) and Xylazine (100 mg/ml).

Following a small midline incision of the skin over the lumbar spine, AAV-RARA, AAV-RARG, or a combination of both were administered at the L3–L6 intervertebral spaces. Briefly, subjects were placed in a prone position to expose the lumbar region. A 10-μL Hamilton syringe was employed to deliver 0.6 μL per site of AAV stock (∼10^11^ viral particles/µL) into the subarachnoid space; a characteristic tail flick served as confirmation of correct needle placement. Post-injection, the musculature was repositioned, the skin was closed with sutures, and the animals recovered on a heating pad. Seven days post-administration, all subjects were euthanized to harvest contralateral lumbar DRGs for primary culture.

### Primary DRG culture

Primary cultures of DRG neurons were established using previously described protocols with minor modifications. Briefly, mice previously injected with AAV-RARA, AAV-RARG, or the combination were sacrificed in accordance with IAEC guidelines. Lumbar DRGs were isolated in pre-warmed Dulbecco’s Modified Eagle Medium/F-12 (DMEM/F-12; Gibco, 12500-062) and digested with Collagenase-I (1 mg/ml; Sigma-Aldrich, SCR103) and 0.05% trypsin (Sigma, #T4799) at 37°C. The enzymatic reaction was halted using DMEM/F12 supplemented with 10% Fetal Bovine Serum (Gibco, #10438026) and 1X penicillin-streptomycin-gentamicin (PSG), followed by mechanical trituration to generate a single-cell suspension. These dissociated cells were seeded onto 96-well plates pre-coated with Poly-D-Lysine (0.1 mg/ml; Gibco, #A3890401) and maintained at 37°C in a 5% CO_2_ environment. After a 48-hour incubation period, the cells were fixed and stained with DAPI for nuclear visualization, followed by the quantification of neurite growth.

### Custom Reference Generation

To detect transgene-expressing nuclei, the AAV-GFP, AAV-RARA and AAV-RARG transgene sequences were appended as separate contigs to the mm10 reference FASTA, and matching transcript and exon entries were added to the GTF annotation. A custom reference was built with cellranger mkref (v7.2). Raw FASTQ reads were aligned to this augmented reference with cellranger count (v7.2; --include-introns=true, --create-bam=true]), producing gene-barcode matrices and coordinate-sorted BAM files. Transgene-positive barcodes were called from the BAM files and count matrix cellular barcodes with ≥1 UMI count for the transgene feature were extracted directly from the Cell Ranger count matrix. These barcodes were added to the Seurat (v5.1) object metadata as a categorical label, enabling separation of transgene-positive and transgene-negative populations for UMAP visualization and differential expression analysis (https://github.com/VenkateshLab/Combinatorial-Control-of-Corticospinal-Axon-Growth-by-Retinoic-Acid-Receptors).

### snRNA-seq analysis

Raw BCL files were demultiplexed and FASTQ reads inspected with FastQC to check base quality and adapter content^23^. Adapters and low-quality bases were removed with fastp v0.23.2^24^, and clean reads were aligned to the mm10 reference using Cell Ranger v7.2 (10x Genomics). Single-nucleus libraries were prepared using the Chromium Next GEM Single Cell 3’ Kit v3.1 [10x Genomics, #PN1000121], sequenced using HiSeq 2500. Gene-barcode matrices were imported into Seurat v5.1 in R^25^, High-quality nuclei were retained by filtering for a minimum of 1,000 features and less than 5% mitochondrial transcripts, and normalized by SCTransform regularized negative binomial regression^26^. Principal component analysis was performed on 30 components with UMAP visualization^27^ yielded 13 transcriptional clusters from 9,891 nuclei (20,174 genes) passing quality control, out of 10,000 nuclei in the raw matrix. Clustering used the Louvain algorithm^28^ on a shared nearest-neighbour graph (9,891 nodes, 246,877 edges; maximum modularity 0.6634 across 10 random starts).

Cell-type identity was assigned by evaluating expression of canonical marker genes across each cluster (Supplementary figure 5), The CST neuron population (n = 6,254), transgene-positive nuclei were identified directly from the custom reference count matrix: 618 nuclei (9.88%) expressed AAV-GFP, 2,221 nuclei (35.51%) expressed AAV-RARA, and 1,661 nuclei (26.56%) expressed AAV-RARG; 1,754 nuclei (28.04%) co-expressed AAV-RARA and AAV-RARG. Rest of the 3,637 as non-CST cell types.

Differential expression between each transduced condition (AAV-RARA, AAV-RARG, AAV-RARA+AAV-RARG and AAV-GFP) was performed with Seurat’s FindMarkers (Wilcoxon rank-sum test, min.pct = 0.05, no log2 fold-change pre-filter). Genes were considered significant at nominal p < 0.05; Benjamini-Hochberg-adjusted p-values are reported alongside nominal p-values in (Supplementary table S6). Gene Ontology biological process enrichment on the resulting differentially expressed gene sets was performed using the DAVID Functional Annotation Tool^29^, separately for genes shared across all three conditions and for genes unique to each condition.

### CUT&RUN data processing

Adapter trimming and low-quality filtering were carried out with fastp v0.23.2^24^. Clean reads were mapped to mm10 using Bowtie2 v2.4.4^30^ with parameters optimized for CUT&RUN fragment recovery (--local --very-sensitive-local --no-mixed --no-discordant --phred33 -I 10 -X 700), converted and sorted with SAMtools v1.13 ^31^. PCR duplicates were marked and removed, using Picard MarkDuplicates. Peaks were called against the shared Input control with MACS2 v2.2.7.1^32^ in paired-end mode (-f BAMPE -g mm -p 1e-4), so every reported binding site is defined as enrichment over the matched background (Supplementary table S8).

Peak-to-gene annotation was performed with the annotatePeak function of ChIPseeker^33^ in R against the mm10 reference, using TxDb.Mmusculus.UCSC.mm10.knownGene for transcript models and org.Mm.eg.db for gene symbol annotation. Each peak was assigned to the gene with the nearest transcription start site. peaks within ±3 kb of a TSS were classified as promoter-associated; peaks outside this window were classified as distal and assigned to their nearest annotated gene by genomic distance. Motif enrichment at RARA and RARG binding sites was assessed against JASPAR motifs [MA0159.1, MA0512.2, MA0730.1, MA0860.1] using HOMER.

Direct transcriptional targets of RARA and RARG were defined by intersecting CUT&RUN peak-annotated genes with the neuronal differential expression gene sets described above. A gene was classified as a putative direct target if it was differentially expressed and had an annotated CUT&RUN peak.

Custom figures UMAPs, heatmaps, bar plots were generated in Python (pandas, NumPy, Matplotlib, SciPy) and R at 600 dpi. All analysis and figure-generation code, parameter files, and processed matrices are available at (https://github.com/VenkateshLab/Combinatorial-Control-of-Corticospinal-Axon-Growth-by-Retinoic-Acid-Receptors).

## Data availability

All genomics datasets generated in this study have been deposited in the NCBI Sequence Read Archive (SRA) under accession number PRJNA1498551. A detailed summary of datasets and corresponding sample information is provided in (Supplementary table S12). The code used for data processing and figure generation is available at (https://github.com/VenkateshLab/Combinatorial-Control-of-Corticospinal-Axon-Growth-by-Retinoic-Acid-Receptors).

## Supplementary Figures

**Supplementary Fig 1** : SDS-PAGE confirms intact capsid formation

**Supplementary Fig 2:** Transgene expression validation for RARA and RARG

**Supplementary Fig 3:** Immunohistochemistry confirms overexpression of the transgenes in AAV-RARA and AAV-RARG treated in-vivo motor cortex samples

**Supplementary Fig 4:** GFAP intensity confirms the extent of spinal cord injury

**Supplementary Figure 5:** Single-nucleus RNA sequencing (snRNA-seq) identification of Corticospinal Tract (CST) and Non-CST neuronal populations

## Supplementary Tables

**Supplementary Table 1.** Curated pro-growth gene list (n = 628) with associated Gene Ontology terms

**Supplementary Table 2.** Neurite length measurements in Neuro-2a cells across AAV treatment groups

**Supplementary Table 3.** Neurite length measurements in primary cortical neurons

**Supplementary Table 4.** Corticospinal axon sprouting quantification after pyramidotomy (fiber index at 200, 400, and 600 µm from midline)

**Supplementary Table 5.** Corticospinal axon regeneration quantification after thoracic crush injury (fiber index at 500, 1000, and 1500 µm from lesion)

**Supplementary Table 6.** Differentially expressed genes from single-nucleus RNA sequencing of the injured motor cortex

**Supplementary Table 7.** Gene Ontology enrichment analysis of single-nucleus RNA sequencing differentially expressed genes

**Supplementary Table 8.** RARA and RARG CUT&RUN peak calls and target gene annotations

**Supplementary Table 9.** Gene Ontology enrichment analysis of RARA and RARG CUT&RUN target genes

**Supplementary Table 10.** Neurite length measurements in dorsal root ganglion neurons

**Supplementary Table 11.** Hindlimb kinematic gait parameters and grip strength measurements after thoracic crush injury

**Supplementary Table 12.** Software, code repositories, and analysis parameters

