## Supplementary figure for "Combinatorial Control of Corticospinal Axon Growth by Retinoic Acid Receptors"

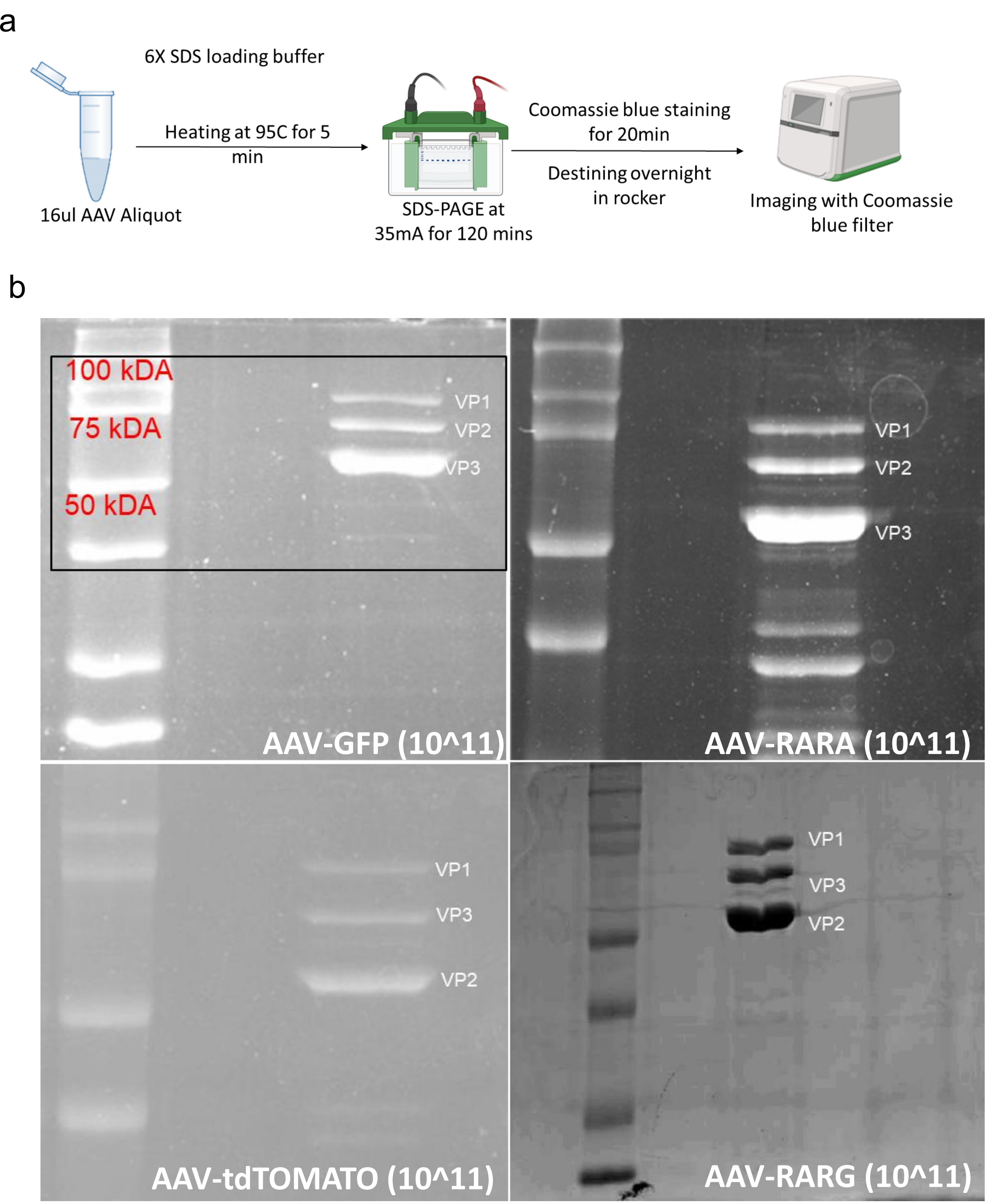


**Supplementary Fig 1 : SDS-PAGE confirms intact capsid formation**

**a.** Schematic of the SDS-PAGE and Coomassie blue staining workflow. AAV aliquots (16 µL) were mixed with 6× SDS loading buffer, heated at 95°C for 5 min, separated by SDS-PAGE at 35 mA for 120 min, stained with Coomassie blue for 20 min, destained overnight, and imaged. **b.** Representative Coomassie-stained gels of AAV-GFP, AAV-RARA, AAV-tdTOMATO, and AAV-RARG preparations at approximately 10^11 viral genomes. The characteristic AAV capsid proteins VP1, VP2, and VP3 were detected at their expected molecular weight ranges, confirming the presence and integrity of the viral capsid protein components.

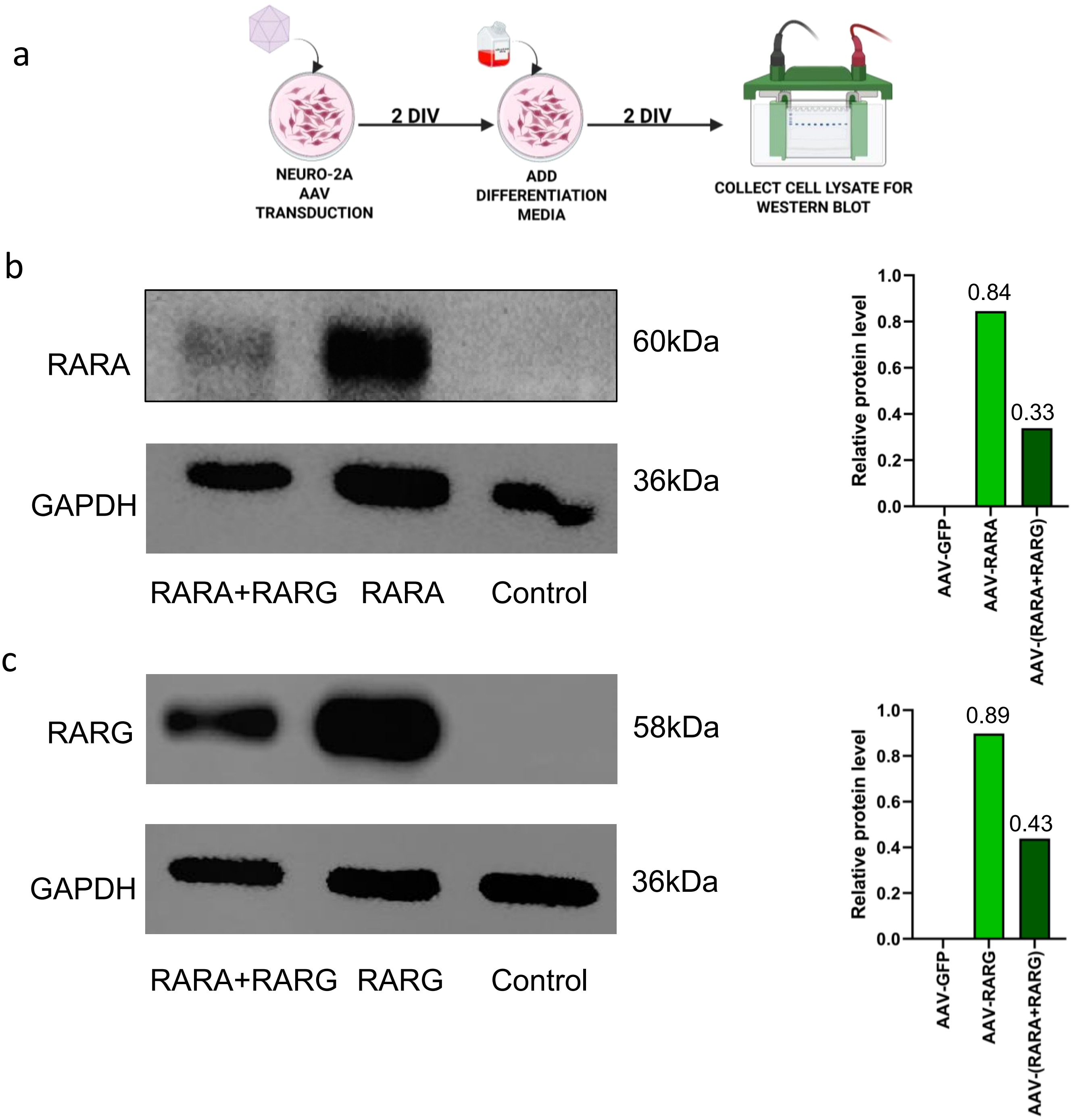


**Supplementary Fig 2: Transgene expression validation for RARA and RARG**

**a.** Schematic illustrating the experimental timeline for Neuro-2A cell transduction and differentiation. Cells were transduced with AAV, followed by the addition of differentiation media for 2 days in vitro (DIV). Cell lysates were collected for Western blot analysis.

**b.** Western blot showing RARA expression in the three conditions: RARA+RARG overexpression, RARA overexpression, and control along with the GAPDH loading controls for the same conditions. The membrane was probed with an RARA antibody and the GAPDH area was probed with GAPDH antibody.

**c.** Western blot showing RARG expression in the three conditions: RARA+RARG overexpression, RARG overexpression, and control along with the GAPDH loading controls for the same conditions. The membrane was probed with an RARG antibody and the GAPDH area was probed with GAPDH antibody.

**d.** Bar plots displaying the normalized expression levels of RARA (top) and RARG (bottom) across the conditions. The expression levels are shown relative to the respective GAPDH loading control of each condition. Data are presented as the mean normalized expression ratio, with each bar representing a single replicate. The numeric values above the bars indicate the specific expression ratios for each condition.

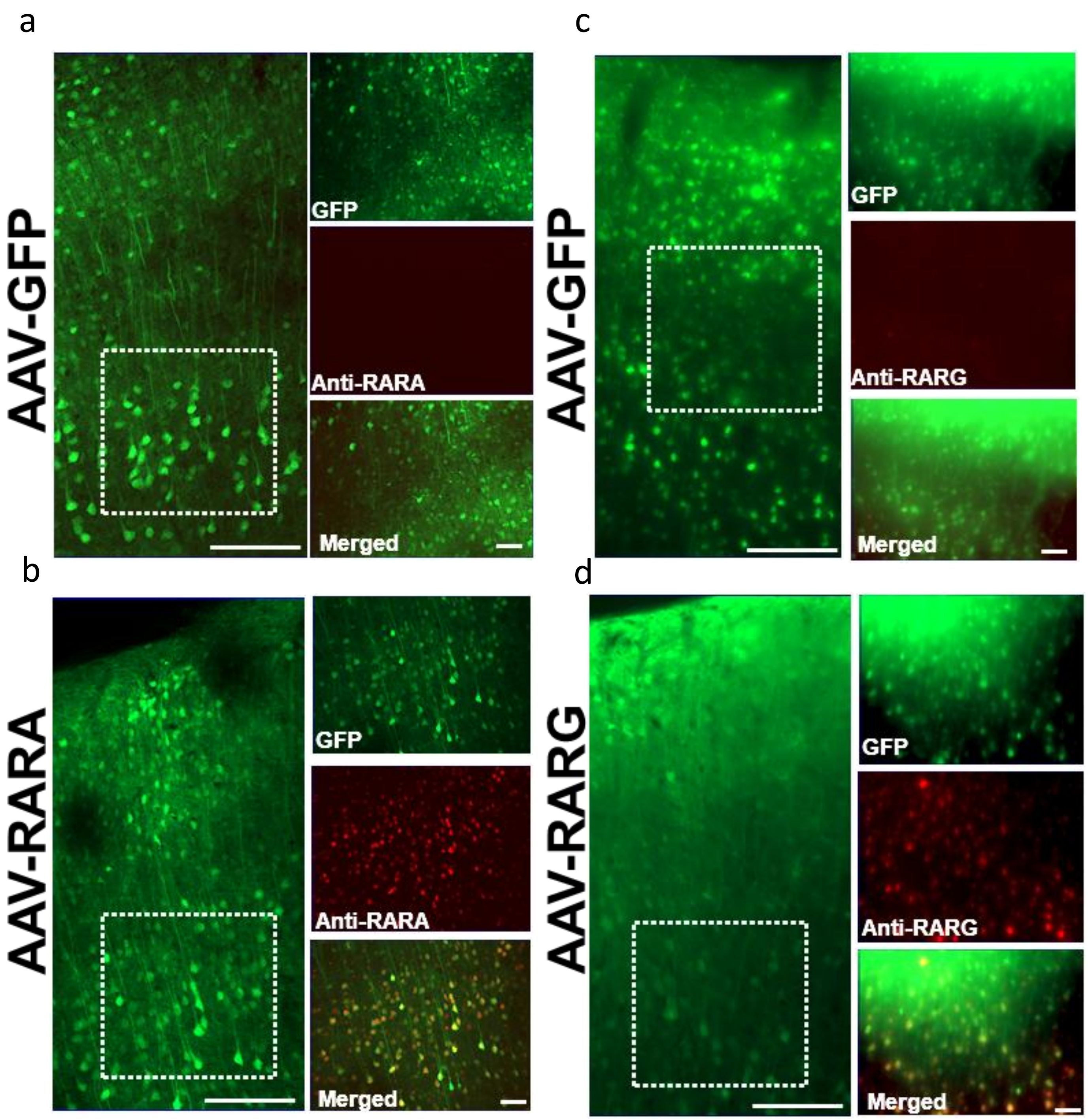


**Supplementary Fig 3: Immunohistochemistry confirms overexpression of the transgenes in AAV-RARA and AAV-RARG treated *in-vivo* motor cortex samples**

Representative fluorescence images of motor cortex sections from animals injected with AAV-GFP, AAV-RARA, or AAV-RARG. **(a–b)** GFP fluorescence and RARA immunoreactivity in AAV-GFP control and AAV-RARA-treated samples, respectively. **(c–d)** GFP fluorescence and RARG immunoreactivity in AAV-GFP control and AAV-RARG-treated samples, respectively. GFP is shown in green, while RARA or RARG immunoreactivity is shown in red. Merged images demonstrate increased RARA and RARG expression in the corresponding AAV-treated groups compared with AAV-GFP controls. Dashed boxes indicate regions shown at higher magnification. Scale bars are 100µm for larger and 50µm for smaller panel.

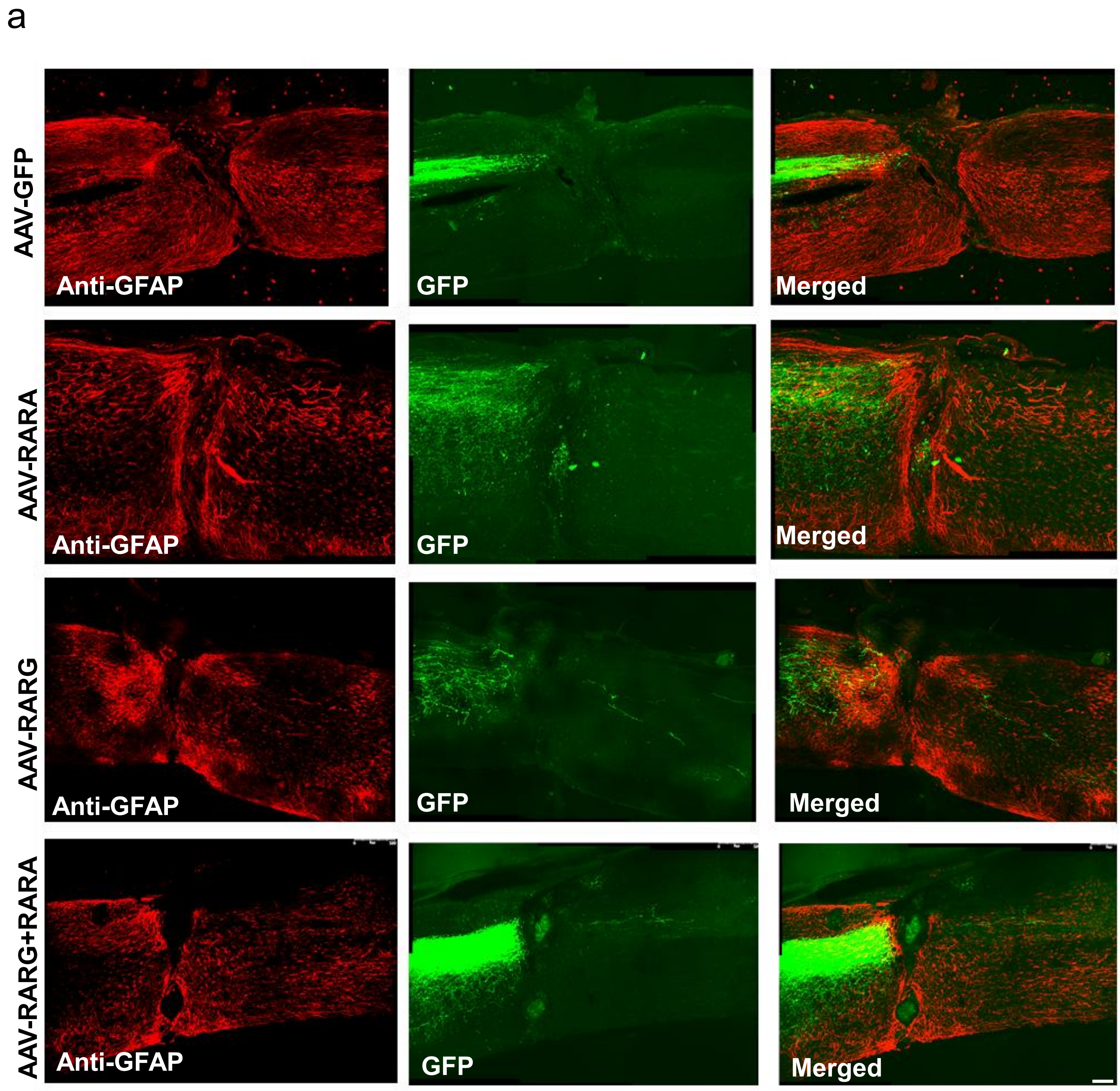


**Supplementary Fig 4: GFAP intensity confirms the extent of spinal cord injury**

**a.** Representative fluorescence images of injured spinal cord sections from animals treated with AAV-GFP, AAV-RARA, AAV-RARG, or combined AAV-RARA+AAV-RARG. GFAP immunoreactivity is shown in red, GFP expression indicating AAV transduction is shown in green, and the corresponding merged images are presented in the right column. GFAP accumulation surrounding the lesion site demonstrates the injury-associated astroglial response. Scale bar, 100 µm.

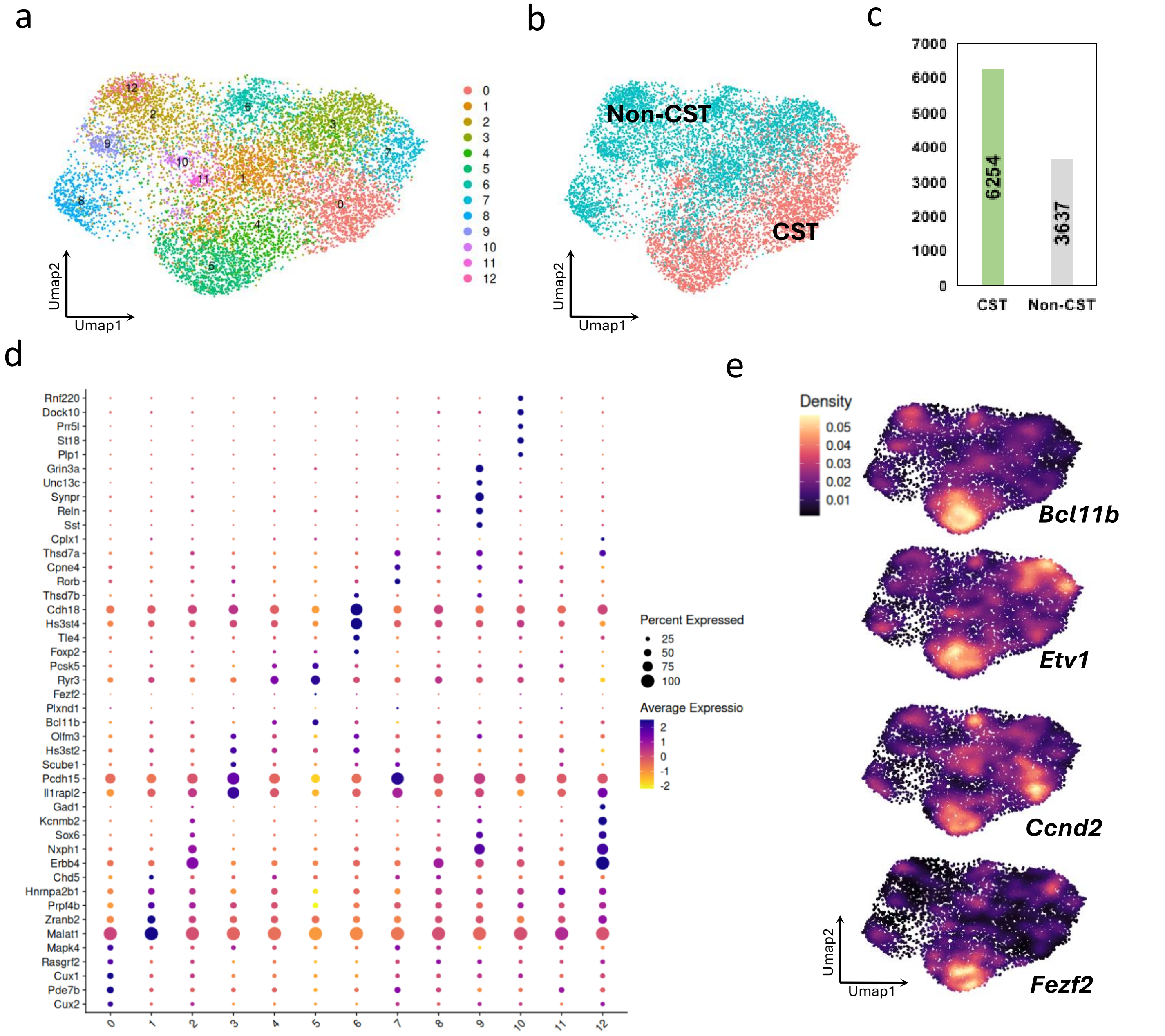


**Supplementary Figure 5. Single-nucleus RNA sequencing (snRNA-seq) identification of Corticospinal Tract (CST) and Non-CST neuronal populations.**

**a.** Uniform Manifold Approximation and Projection (UMAP) embedding of snRNA-seq data, revealing 13 distinct transcriptional clusters (0-12). **b.** UMAP projection with clusters annotated and grouped into Corticospinal Tract (CST, red) and Non-CST (teal) populations. **c.** Bar plot quantifying the total number of recovered nuclei assigned to the CST (n = 6,254) and Non-CST (n = 3,637) lineages. **d.** Dot plot illustrating the expression profiles of selected marker genes across all 13 clusters. Dot size represents the percentage of cells expressing the gene within a specific cluster, and the color gradient indicates the average scaled expression level. **e.** Feature density plots visualizing the localized expression of canonical CST marker genes (Bcl11b, Etv1, Ccnd2, and Fezf2) across the UMAP embedding.
